# Data-driven model reveals increased stability of CAG-expanded *huntingtin* RNA due to MID1 binding

**DOI:** 10.1101/2025.10.31.685889

**Authors:** Yuhong Liu, Annika Reisbitzer, Domagoj Dorešić, Jan Hasenauer, Sybille Krauß, Tatjana Tchumatchenko

**Author notes:** These authors contributed equally (Jan Hasenauer, Sybille Krauß, Tatjana Tchumatchenko).

## Abstract

RNA-binding proteins (RBP) are important regulators of RNA metabolism. In neurodegenerative disorders such as Huntington’s Disease (HD), disrupted RBP-RNA interactions contribute to neuronal dysfunction. One such RBP, Midline 1 (MID1), has been shown to aberrantly associate with mutant huntingtin (*Htt*) RNA, enhancing its translation, yet the mechanism driving this effect remains unknown. Here, we develop a computational model to understand the role of MID1. Based on previously published data, our model predicts that MID1 increases the stability of the *Htt* RNA. We experimentally validate this prediction, showing that overexpression of MID1 significantly prolongs the half-life of mutant *Htt* RNA. Furthermore, we evaluate model refinements, including clustering of MID1-bound RNA, which allow capturing all key observations in the data. Together, we provide a data-driven framework that underlines the importance of RBP-RNA interaction in post-transcriptional regulation. This framework also shows how individual molecular reactions jointly determine RNA stability and protein levels in HD.

## Introduction

Translation of RNA into protein is a fundamental cellular process that influences the amount of proteins available for cellular activity. ^5,55^ Such a process is subject to multilayered regulation, both temporally^6^ and spatially, ^37,10,45^ and even minor perturbations can result in severe cellular dysfunction. ^25,11,23^ A central mechanism for this regulation involves interactions between RNA and RNA-binding proteins (RBP), which coordinate key steps of RNA metabolism such as splicing, localization, stability, and translation efficiency. ^18,42,14^ Disruptions in these interactions are particularly detrimental in neurons, where RNA processing plays a pivotal role in maintaining neuronal integrity and function. ^47,33^ Indeed, aberrant RBP activity is increasingly recognized as a pathogenic driver in various neurological disorders.^34,54^

A prominent example is Huntington’s Disease (HD), a neurodegenerative disorder caused by an expansion of the CAG trinucleotide repeats in the huntingtin gene (*Htt*).^31,32^ The CAG trinucleotide encodes glutamine (Q) in the huntingtin protein (HTT), so, in the following, we use Q number and CAG repeat length interchangeably. The number of CAG repeats in *Htt* RNA is typically fewer than 36 in the wild-type (wt).^52^ Mutant *Htt* RNA, however, contains longer CAG repeats, which fold into stable hairpin structures. ^27,15^ These aberrant secondary structures found in *Htt* RNA with expanded CAG repeat lengths are preferentially recognized by the RBP Midline 1 (MID1) (Fig. 1A, known interaction 1), an E3 ubiquitin ligase tagging proteins for degradation.^46,48^ This specific interaction between MID1 and mutant *Htt* RNA can thus potentially modify RNA metabolism.

**Figure 1.**
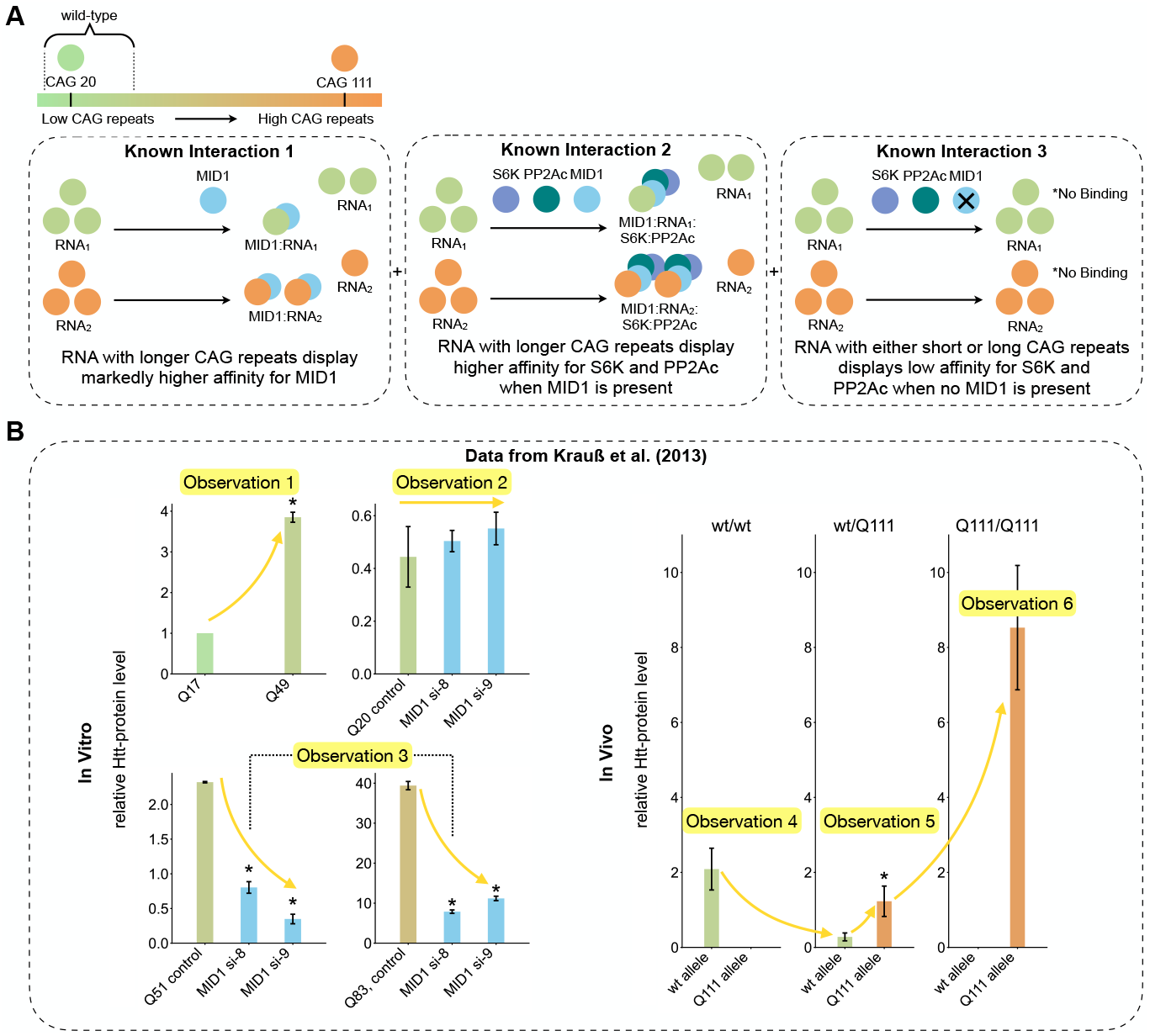
Visual summary of impact of CAG-expansion, and the experimental data used for parameter inference. (A) Schematic representation of MID1-dependent RNP complex formation. The legend indicates the color scheme used to represent different CAG repeat or glutamine (Q) lengths. Wild-type is defined as when the CAG repeat or glutamine length is less than 36. (B) Experimental *in vitro* and *in vivo* data taken from^27^. In the *in vitro* panels, the x-axis labels, e.g., Q17, indicate constructs with the corresponding number of CAG repeats, while in the *in vivo* panels, the subtitles, e.g., wt/wt, denote mouse genotypes. MID1 si-8 and MID1 si-9 refer to two independent small interfering RNA sequences specifically designed to knock down MID1 expression. The error bars represent the standard deviation of the data. An asterisk above a bar indicates a significant difference (*p <* 0.05) relative to the first bar in the subplot.

Upon binding to *Htt* RNA, MID1 orchestrates the assembly of a translationally active ribonucleoprotein (RNP) complex that includes protein phosphatase 2A (PP2A) and S6 kinase (S6K).^15,2^ This interaction also occurs in a CAG repeat length-dependent manner:^27^ RNA with longer CAG repeat length (e.g., 111 CAG repeats) displays markedly higher affinity for MID1 and its associated complex compared to those with physiological repeat lengths (e.g., 20 CAG repeats) (Fig. 1A, known interaction 2). The binding affinity is length-dependent because the CAG-containing RNA hairpin progressively elongates and stabilizes as the CAG repeat length increases, thereby providing a larger binding platform for MID1.^27^ Notably, knockdown of MID1 abolishes PP2A and S6K association with *Htt* RNA, establishing MID1 as a necessary scaffold for complex formation (Fig. 1A, known interaction 3). ^27^

These recruited complex compounds, PP2A and S6K, modulate phosphorylation states of translation regulators, thereby enhancing translation initiation of the mutant RNA. The downstream consequence of these interactions is an increase in translation efficiency from RNA with expanded repeats. This is evident in published data for *in vitro* and *in vivo* experiments (Fig. 1B).^27^ In *in vitro* experiments, translation of *Htt* RNA constructs bearing high CAG repeat length yields substantially more protein (e.g., Q49, which indicates HTT with a polyglutamine stretch of 49 residues) than constructs with lower CAG repeat length (e.g., Q17), despite similar RNA levels. This effect is abolished upon MID1 knockdown using MID1-specific short interfering RNA (MID1 si-8 or si-9) for constructs with expanded repeats, while having minimal or no effect on low-repeat constructs—likely due to the weak binding affinity between MID1 and wt *Htt* RNA.^27^

Furthermore, additional *in vivo* evidence from knock-in mouse models carrying the mutant *Htt* allele (Q111/+) suggests that simple binding interactions might not be sufficient to explain the observed data (Fig. 1B *in vivo* panel). In both heterozygous and homozygous mutant mice, the mutant allele consistently produces higher levels of HTT than the wt allele, mirroring the *in vitro* findings. However, the data also reveals more complex protein level patterns that cannot be explained by a simple linear increase in MID1 binding. For example, in heterozygous mice, while the mutant allele shows significantly higher protein output than the wt allele as expected, the total amount of protein is comparable with that in homozygous wt mice. This suggests that MID1 binding alone does not fully account for the differences. Moreover, although homozygous mice carry twice as many mutant alleles as heterozygous mice, their HTT levels are more than double, indicating a supralinear increase that cannot be explained by allele dosage alone.

These experimental findings together demonstrate that MID1 increases the translation of RNA with expanded CAG repeats. While the role of MID1 in recruiting translation-associated kinases is well established, the quantitative link between these molecular interactions and the resulting differences in protein expression remains unresolved. In particular, it is still unclear how the difference in CAG repeat length, MID1 binding affinity, and co-factor recruitment jointly determine translation efficiency and drive the observed expression patterns. Furthermore, these observations indicate that other mechanisms introducing nonlinear dependencies, such as competitive or cooperative effects, may be at play beyond the simple binding affinity or allele dosage.

Mathematical modeling is a powerful approach to unravel such mechanisms. Ordinary differential equation (ODE) models have already proven to be powerful tools for elucidating the quantitative dynamics of RNA–protein interactions,^19,35^ as well as autoregulatory feedback loops of RBP.^8^ Furthermore, ODE models have been used to investigate the role of the yeast RBP Puf3 in regulating RNA stability. ^24^ However, despite these advances, such quantitative frameworks have yet to be extensively applied to understand RBP-driven RNA translation dynamics in neurodegenerative diseases like HD.

Our work addresses this gap by developing an ODE model describing the influence of CAG repeat length on MID1 binding and the resulting HTT levels based on the data^27^ described above (Fig. 1B). We derive testable predictions, such as the impact of MID1 binding on *Htt* RNA stability. To evaluate these predictions, we conduct biological experiments. Furthermore, we use parameter estimation and model selection to assess which model best describes the available data. By integrating mathematical modeling with experimental data, our findings provide a new mechanistic understanding of how MID1 influences RNA regulation and offer potential targets for therapeutic intervention in disorders characterized by abnormal RNA stability and translation dynamics.

## Results

### MID1-dependent core molecular interactions alone do not account for the observed protein level patterns

To systematically uncover how MID1 influences HTT level, we consider previously published experimental data regarding several characteristic observations of HTT level in the presence of expanded CAG repeats (Fig. 1B).^27^ These observations are:

(Observation 1)HTT levels increase with the CAG repeat length.

(Observation 2)MID1 level has a minimal effect at low CAG repeat lengths.

(Observation 3)MID1 level modulates HTT levels at high CAG repeat lengths.

(Observation 4)The wt allele produces a lower HTT level in wt/Q111 compared to wt/wt mice.

(Observation 5)In wt/Q111 mice, the Q111 allele is preferentially translated.

(Observation 6)Q111/Q111 mice show a supralinear increase in the HTT level from the Q111 allele in wt/Q111 mice.

To evaluate whether these characteristics can be explained solely by the core molecular interactions between *Htt* RNA, MID1, and downstream effectors, we develop a *Baseline Model*. This model captures the previously identified core molecular interactions (Fig. 1A), including allele-specific MID1 binding, MID1-dependent recruitment of the S6K, phosphorylation-dependent translation, and protein degradation.

The *Baseline Model* considers two *Htt* RNAs (one from each allele; denoted RNA_1_, RNA_2_), unbound MID1, MID1-bound RNA complexes (MID1:RNA_1_, MID1:RNA_2_), unphosphorylated and phosphorylated S6K (S6K, S6K^P^), and the resulting HTT (HTT_1_, HTT_2_). The concentrations of all biochemical species in the model are encoded in the state vector:

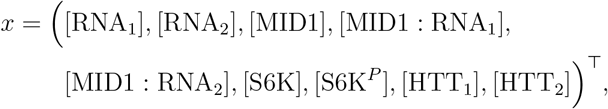

The characteristics of RNA_1_ and RNA_2_, i.e., CAG repeat lengths *q* of each allele, *q*_1_ and *q*_2_, depend on the specific experimental context. For instance, for an experiment with heterozygous wt/Q111 mice, the RNA_1_ is the wt RNA and RNA_2_ is the Q111 RNA. As the model structure is symmetric with respect to RNA_1_ and RNA_2_, the order of the assignment will not influence results (see Supplementary Information Sec. S1.1).

The model accounts for four classes of core reactions, which are illustrated in SBGN^4,38^ (Fig. 2A):

**Figure 2.**
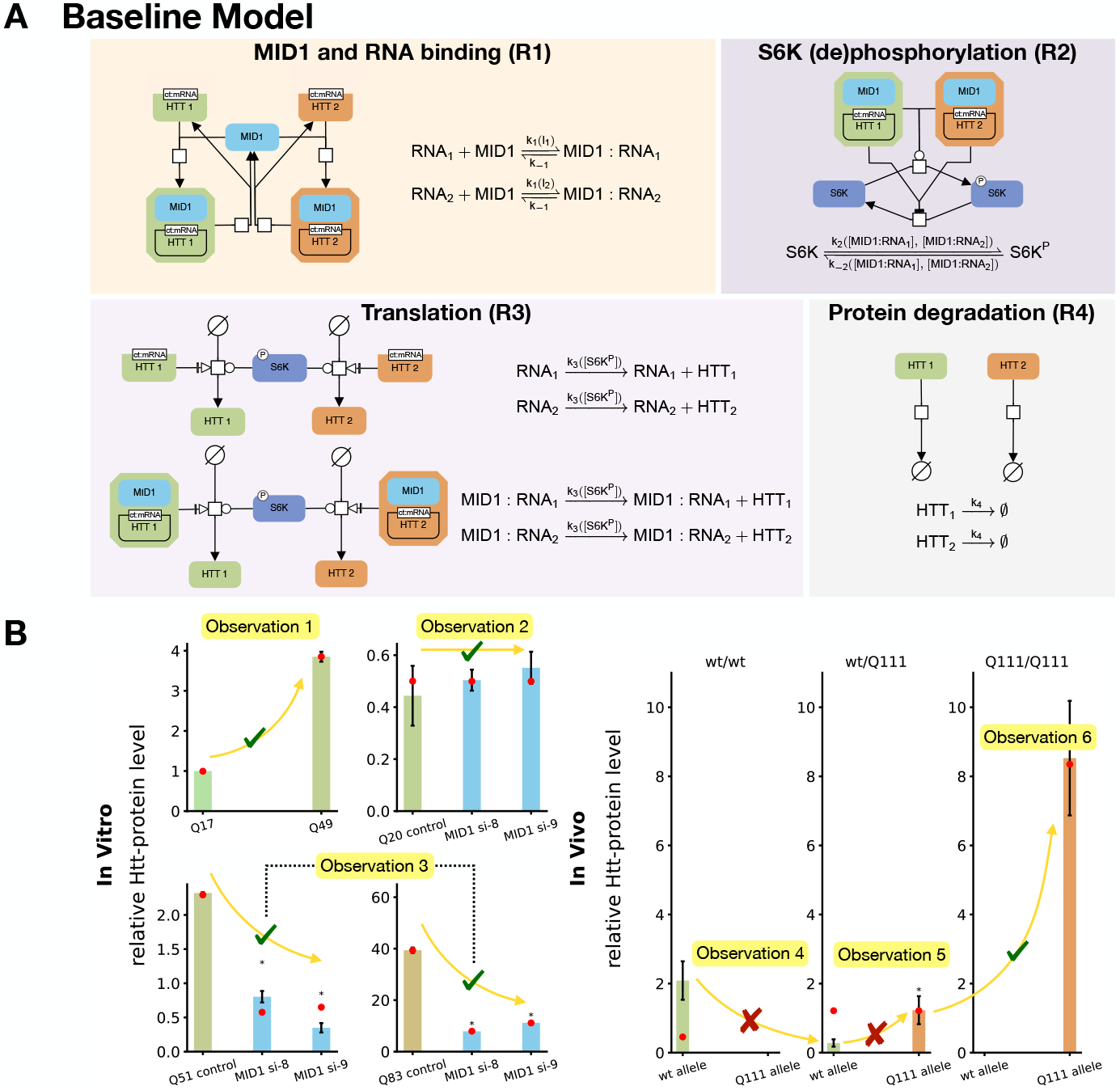
*Baseline Model*’s SBGN visualization and fitting results. (A) Visualization of the *Baseline Model* using the Systems Biology Graphical Notation (SBGN) language, showing the binding of MID1 and RNA has a downstream effect on S6K phosphorylation and subsequently translation. (B) Simulation results of the *Baseline Model* for the maximum likelihood estimate (red dot): only four out of six data observations are reproduced. The error bars represent the standard deviation of the data. An asterisk above a bar indicates a significant difference (*p <* 0.05) relative to the first bar in the subplot.

#### (R1) CAG length-dependent binding of MID1 to *Htt* RNA

*Htt* RNA binds MID1 in a manner dependent on the CAG repeat length *q* of each allele, i.e., *q*_1_, *q*_2_. We implement this dependence upon *q* via a function *l*(*q*), which is continuously increasing with respect to *q* (see Supplementary Information Sec. S4), following mass action kinetics:

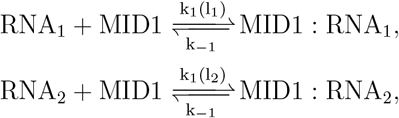

where the forward rate is proportional to *l*, e.g., *k*_1_(*l*_1_) = *c*_1_ · *l*_1_(*q*_1_), and the unbinding rate *k*_−1_ = *c*_−1_ is independent of CAG repeat length. In the following text, *k* represents the reaction rate and *c* represents the reaction rate constant.

#### (R2) S6K phosphorylation modulated by MID1-bound RNA

Binding of MID1 to *Htt* RNA promotes S6K phosphorylation and inhibits its dephosphorylation:

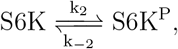

with the reaction rates:

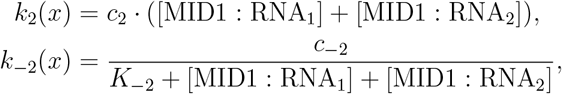

where *x* is the state vector and *K*^−2^ is the inhibition constant for MID1:RNA mediated suppression of S6K dephosphorylation. Note that the [S6K^P^] modeled here only represents the concentration level modulated by MID1. We discuss how phosphorylated S6K, modulated by MID1 or not, affects translation rate in the following reaction (R3).

#### (R3) Translation as a function of S6K^P^ abundance

Translation occurs for both free and MID1-bound RNA and is positively regulated by phosphorylated S6K modulated by MID1:

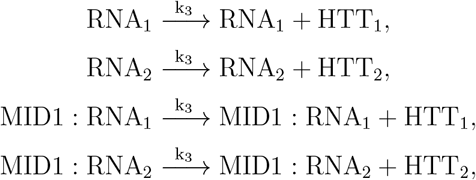

with the reaction rates:

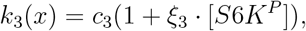

where *c*_3_ is the basal translation rate that includes all other sources of translational activity in the cell. For instance, *c*_3_ includes all other effects on the translation rate from phosphorylated S6K not directly modulated by MID1. We assume the effect from the S6K^P^ modulated by MID1, *c*_3_ · *ξ*_3_ · [S6K^P^], cannot be too large compared to the basal translation and constrain *ξ*_3_ to values between 0 and 100.

#### (R4) Degradation of HTT

The fourth reaction is first-order degradation of HTT:

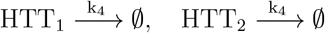

with the degradation rate *k*_4_ = *c*_4_.

The list of reactions was translated into an ODE model. The system of ODEs governing the dynamics of the *Baseline Model* is provided in the Supplementary Information Sec. S1.1. This model and all subsequent models admit a unique solution that remains non-negative for all *t* ≥ 0 (see Supplementary Information Sec. S5). The initial RNA levels are assumed to be the same and are estimated using the parameter *rna*_0_. The initial MID1 level depends on specific experiment conditions, i.e., on the presence of siRNAs (see Supplementary Information Sec. S2). The initial S6K level is estimated using the parameter *s*6*k*_0_. All other initial conditions are assumed to be zero.

The CAG repeat length *q* is generally known for most experimental conditions. For instance, the experimental condition Q111/Q111 means that both alleles carry 111 CAG repeats, i.e., *q*_1_ = *q*_2_ = 111. However, for wt/Q111 and wt/wt settings, the CAG repeat length for the wt allele is not exactly known. We thus define the CAG repeat length of the wt allele in wt/Q111 and wt/wt settings to be an estimated parameter, *Q*_*wt*_ (see Supplementary Information Sec. S9). The reaction rate constants, e.g.,*c*_1_, are also unknown parameters.

To test whether the model can explain the data (Fig. 1B), the model parameters are estimated by maximum likelihood estimation. As the data provides information about the HTT abundance in steady state, we have the observables

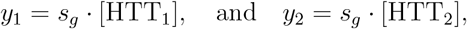

with western blot gel-specific scaling factors *s*_*g*_, *g* ∈ *{*1, 2, …, 5*}*, capturing difference in detection and normalization effects.

Parameter estimation using a state-of-the-art pipeline provides reproducible results (see Supplementary Information Sec. S12). The comparison of data and model simulations reveals that four observations are captured (Fig. 2B), i.e., Observation 1, Observation 2, Observation 3, and Observation 6. However, the model fails to explain two other essential observations of the data, i.e., Observation 4 and Observation 5. These discrepancies suggest that additional molecular mechanisms—beyond the core MID1-dependent molecular interactions encoded in the *Baseline Model* —are required to fully capture the observed protein level patterns.

### Model-based analysis indicates different degradation rates for free and MID1-bound *Htt* RNA

To obtain a more consistent description of the experimental data, we adjust the *Baseline Model* with additional reactions. The *Baseline Model* assumes a constant translatable RNA resource, i.e., *rna*_0_. For a dynamical system with many interacting species, this simplified assumption is, however, rarely holds. Thus, instead of assuming a fixed total *Htt* RNA level for both alleles, we now explicitly model *Htt* RNA synthesis and degradation. The *Extended Model* retains the molecular species and interactions of the *Baseline Model* but introduces three additional reactions (Fig. 3A):

**Figure 3.**
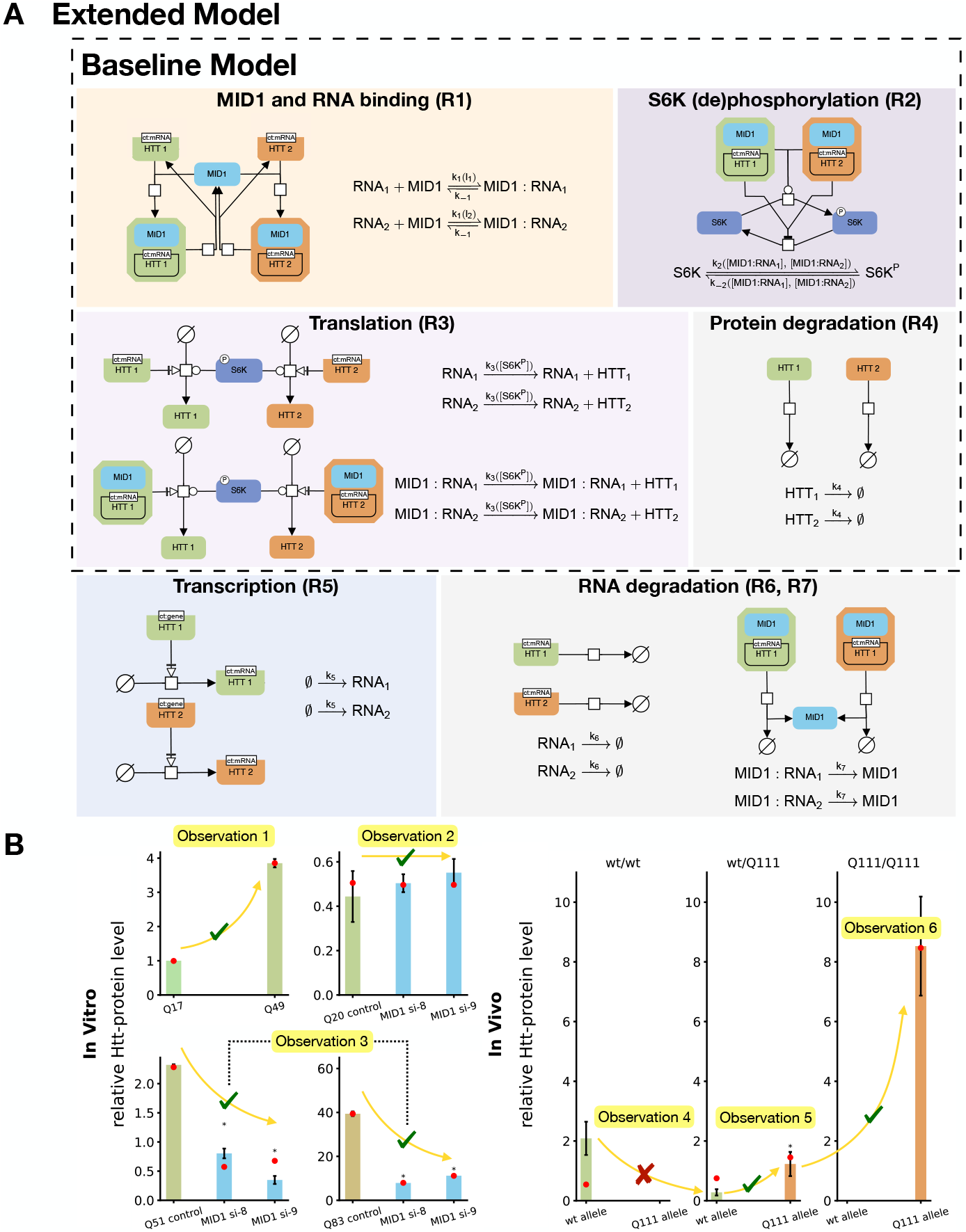
*Extended Model*’s SBGN illustration and fitting results. (A) *Extended Model* builds on *Baseline Model* by incorporating *Htt* RNA synthesis and degradation. (B) The model captures additional Observation 5 compared to the *Baseline Model*, including higher translation of the Q111 allele in heterozygous mice. The error bars represent the standard deviation of the data. An asterisk above a bar indicates a significant difference (*p <* 0.05) relative to the first bar in the subplot.

#### (R5) Transcription of *Htt* RNA

New RNA for both alleles is assumed to be transcribed at a constant rate:

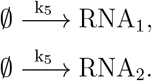

The transcription rate for both alleles is given by *k*_5_ = *c*_5_.

#### (R6) Degradation of free *Htt* RNA

Free *Htt* RNA degrades at a constant rate:

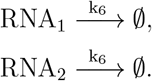

The degradation rate for both alleles is given by *k*_6_ = *c*_6_.

#### (R7) Degradation of MID1-bound *Htt* RNA

MID1-bound *Htt* RNA degrade independently of free *Htt* RNA and release MID1:

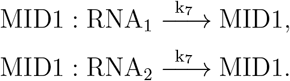

The degradation rate for both alleles is given by *k*_7_ = *c*_7_.

The model formulation does not include a direct dependency of the degradation rates on the CAG repeat length. However, since repeat length affects MID1 binding affinity, and thus the balance between free and bound RNA, an indirect effect on RNA stability can still arise.

The initial condition, states, parameter set, and system of ODE governing the dynamics of the extended model are provided in the Supplementary Information. The parameters of the model incorporating *Htt* RNA synthesis and degradation are determined using maximum likelihood estimation. The same setup as for the *Baseline Model* is applied, and the optimizer outputs indicate the optimization has converged (see Supplementary Information Sec. S12).

The analysis of the estimation results reveals that the *Extended Model* reproduces another important experimental observation compared to the *Baseline Model*. In particular, it correctly recapitulates Observation 5 in addition to the Observations captured by the *Baseline Model* (Fig. 3B). However, the model still fails to reproduce Observation 4, where the wt allele produces a lower HTT level in wt/Q111 mice compared to in wt/wt mice.

Despite its limitations, the *Extended Model* now successfully fits Observation 5 and provides a mechanistic explanation for the higher Q111 protein level compared to the wt protein level in wt/Q111 mice. We analyze the steady state total RNA concentrations (free and MID1-bound) for both alleles and find a consistently higher total RNA level for the Q111 allele (see Supplementary Information Sec. S7). This asymmetry in RNA abundance, despite equal transcription rates, indicates that differences in degradation rates play a central role.

Indeed, our analysis of the inferred parameters shows that the two degradation processes (Fig. 4A) have different rate constants. The degradation rate constant of MID1-bound RNA (*c*_7_) is lower than that of free RNA (*c*_6_) (Fig. 4B). This finding implies that MID1 binding not only enhances translation efficiency via S6K phosphorylation but also stabilizes *Htt* RNA by protecting it from degradation.

**Figure 4.**
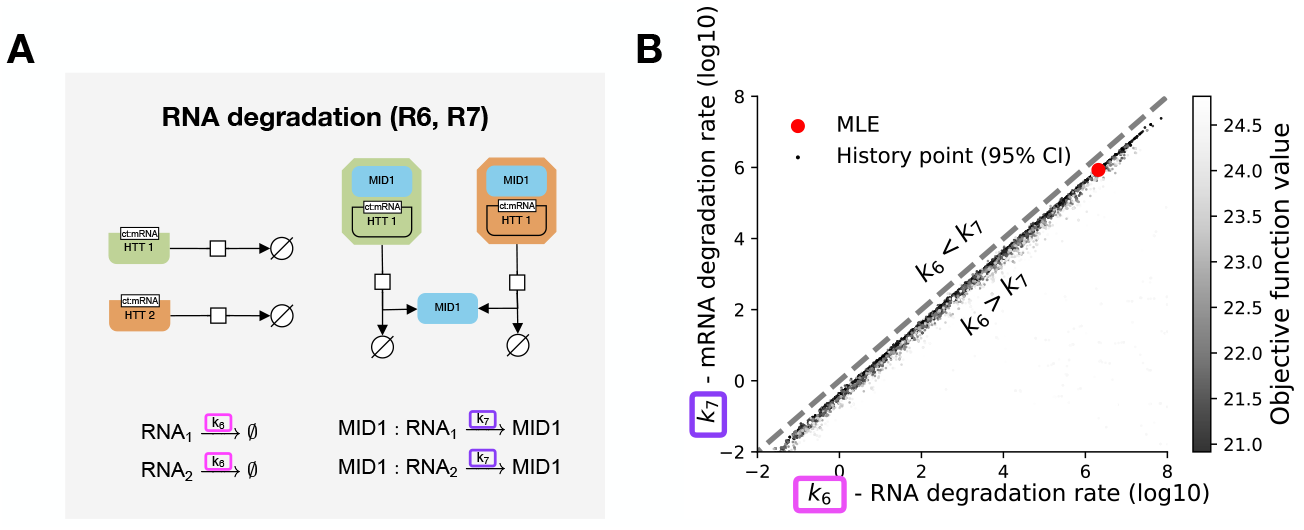
MID1 binding increases RNA stability. (A) Illustration of free and MID1-bound RNA degradation pathways. (B) Scatter plot of (*k*_6_, *k*_7_) parameter pairs (log10 scale). The maximum likelihood estimate (red) and points along the optimizer path (gray scale values according to the objective function) are shown. Only parameter vectors within the 95% confidence region, as defined via the *χ*^2^ threshold, are included.

### Experimental data validates MID1 mediated protection of *Htt* RNA against degradation

To validate the model-based prediction of an altered degradation rate, we conduct a new experiment and assess how *Htt* RNA degrades under two conditions: a control condition (without MID1 transfection) and a MID1 overexpression condition (Fig. 5A). After the establishment of the cell culture, including the induction of a *Htt* allele with 83 CAG repeats (Q83) in the cell and transfection with plasmids coding for MID1, cells are treated with Actinomycin D to inhibit transcription. Subsequently, the relative RNA levels are monitored using qPCR.

**Figure 5.**
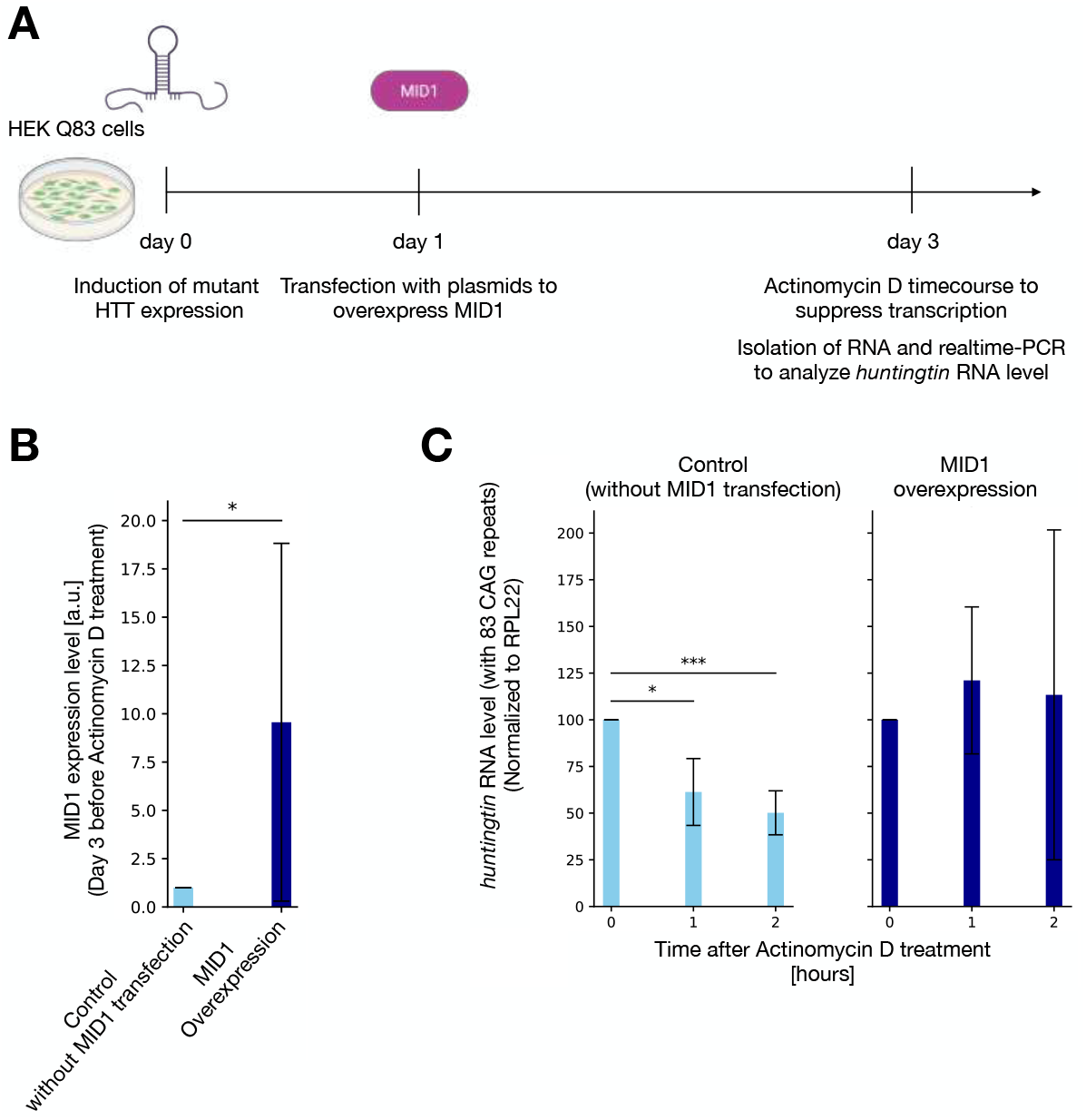
Experimental data verifies the model prediction that MID1 increases RNA stability and thereby the abundance of *Htt* RNA relative to wt. (A) Experimental setup. (B) MID1 level comparison under control (without MID1 transfection) and MID1 overexpression conditions on day 3. (C) Relative *Htt* RNA (with 83 CAG repeats) level within 2 hours under control (without MID1 transfection) and MID1 overexpression conditions. The error bars represent the standard deviation of the data. An asterisk above a bar indicates a significant difference (*p <* 0.05) relative to the first bar in the subplot.

The assessment of the MID1 levels three days after plasmid transfection confirms a substantial increase compared to the control condition (Fig. 5B). The difference in MID1 levels results in different *Htt* RNA decay dynamics (Fig. 5C). Under the control condition (without MID1 transfection), the *Htt* RNA level is reduced by 50 % after two hours, indicating a half-life time of approximately 2 hours. Under the MID1 overexpression condition, the *Htt* RNA levels are stable over the same time interval, implying a substantially longer half-life.

The experimental data provides a validation of our model-based predictions, illustrating the significant functional role of MID1 in regulating RNA stability.

### Clustering and local signaling explain observed HTT levels

The validation experiments show that the *Extended Model* can provide valuable, testable hypotheses. Yet, the model does not recapitulate Observation 4. As the underlying cause of this limitation is unclear, we employ model-based hypothesis testing.

We put a constraint on the *Extended Model* which enforces the experimentally observed increased stability of MID1-bound RNA, i.e., *k*_7_ ≤ *k*_6_ as a starting point for the model-based hypothesis testing. To account for the constraint, we reformulate the degradation reaction as follows:

#### (R6_R_) Degradation of free *Htt* RNA

Free *Htt* RNA degrades at a constant rate:

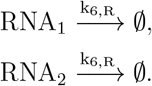

The degradation rate for both alleles is given by *k*_6,*R*_ = *ξ*_6_*k*_7_, where 1 ≤ *ξ*_6_. The parameter *ξ*_6_ encodes the stability difference between free and MID1-bound RNA.

In the following part, we explore three models developed upon the *Extended Model*, including additional mechanisms that may affect the HTT level.

The first possible explanation for the observations might be a nonlinear translation kinetic. As translation is a multi-step process, it might depend nonlinearly on the abundance of S6K^P^. We assess this by considering a Hill-type model, which allows the model to contain a threshold-dependent event, where translation efficiency increases slowly at low activator concentrations but rises sharply beyond a certain threshold. The *Extended Model with Nonlinear Translation* retains the molecular species and interactions from the *Extended Model* but utilizes a Hill-type reaction rate for reaction (R3):

#### (R3_H_) Translation as a function of S6K^P^ abundance

Translation occurs for both free and MID1-bound RNA and is positively regulated by phosphorylated S6K:

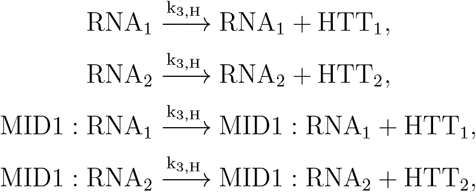

with the reaction rates:

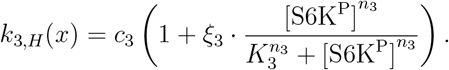

The second possible explanation for the observations is that RNA translation is regulated within spatially restricted subcellular domains, such that S6K^P^ exerts a local rather than global effect. Multiple experimental studies have shown that cytoplasmic RNA is often non-uniformly distributed, becoming enriched in discrete sites such as membraneless condensates,^29^ TIS granules,^20^ or translation factories, ^36^ where local translation rates could differ markedly from those in the surrounding cytosol. A spatial enrichment can arise through a variety of mechanisms, including physical clustering of RNA (often associated with RNA granules),^26,29^ targeted localization to translation-permissive compartments, ^20^ or co-translational recruitment of related RNA through nascent chain interactions.^49,44^ Previous models assume a spatially homogeneous, well-mixed cytoplasm in which molecular species interact uniformly, but this assumption may oversimplify the heterogeneous organization of RNA translation regulation. To address this shortcoming, we model “clusters” in a generalized sense as any localized subpopulation of MID1-bound RNA—regardless of the precise mechanism underlying their enrichment—whose translation can be selectively modulated by locally elevated S6K^P^ activity. Such compartmentalized regulation enables precise, context-dependent protein synthesis and may account for the nonlinear dynamics and allele-specific effects observed in the experimental data. The *Extended Model with Clustering* incorporates spatial structure implicitly through localized translation regulation. It assumes that MID1-bound *Htt* RNA can aggregate into spatially restricted molecular clusters, where they exert local influence on S6K phosphorylation inside the cluster. This localized S6K activation selectively enhances translation of clustered RNA without affecting global translation rates. Outside the cluster, translation is assumed to proceed as in the *Extended Model*. The *Extended Model with Clustering* introduces several new reactions (Fig. 6), and expands the state vector to include the cluster-localized species [MID1 : RNA_1,c_], [MID1 : RNA_2,c_], [S6K_c_], and 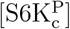.

**Figure 6.**
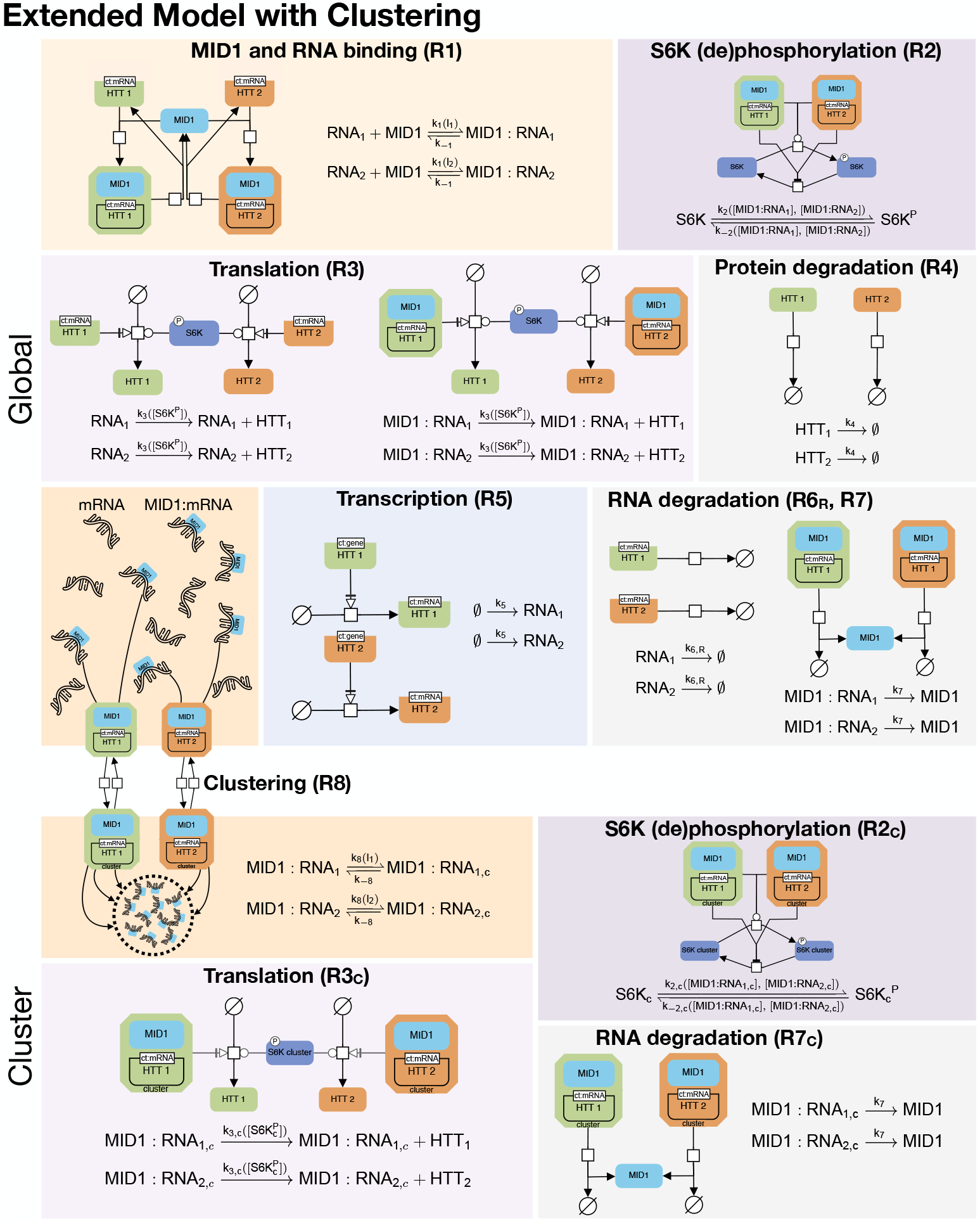
SBGN illustration of the *Extended Model with Clustering*. MID1:RNA complexes form a local cluster that selectively modulates nearby S6K phosphorylation and translation activity.

The additional and modified reactions are:

#### (R8) Cluster association and dissociation

MID1-bound RNA can reversibly join clusters:

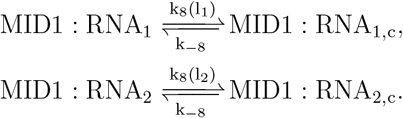

We consider a CAG repeat length dependent association rate for cluster entry, i.e., *k*_8_(*l*) = *c*_8_ · *l*, and a constant dissociation rate *k*_−8_ = *c*_−8_. In the Supplementary Information Sec. S6.1, we show that models lacking CAG repeat length dependence (i.e., *k*_8_(*l*) ≡ *c*_8_) fail to recapitulate the observations, demonstrating that *l* dependence is necessary.

#### (R2_C_) Local S6K phosphorylation

Within the cluster, S6K is phosphorylated in proportion to the amount of MID1-bound RNA present:

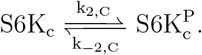

The rates are assumed to be identical to the rates outside the cluster,

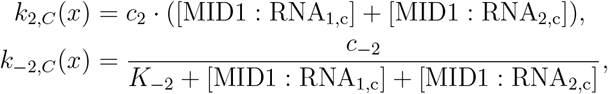

but the effective concentrations differ. The S6K phosphorylation outside the cluster remains the same as in the *Baseline Model* and in the *Extended Model*.

#### (R3_C_) Translation inside clusters as a function of S6K^P^ abundance

MID1:RNA complexes in the clusters are translated in an S6K^P^-dependent manner,

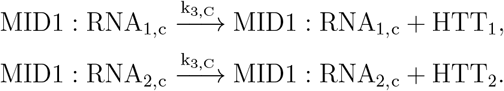

with rate:

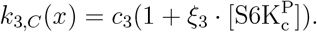

The translation outside the cluster remains the same as in *Baseline Model* and in the *Extended Model*.

#### (R7_C_) Degradation of the MID1:RNA in the cluster

Both global and clusterassociated MID1:RNA species degrade at the same MID1-dependent rate *k*_7_ = *c*_7_, distinct from the degradation of unbound RNA.

As the two mechanisms, nonlinear translation and clustering, are not exclusive, we also consider the *Extended Model with Clustering and Nonlinear Translation*. The model is based on *Extended Model with Clustering*, with the translation outside the clusters changed to (R3_H_) and the translation inside is defined as the following:

#### (R3_C, H_) Translation inside clusters as a function of S6K^P^ abundance

MID1:RNA complexes in the clusters are translated in an S6K^P^-dependent manner,

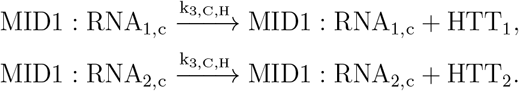

with rate:

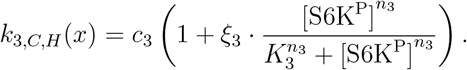

The ODE systems governing the dynamics of all three models are provided in the Supplementary Information Sec. S1. All models are conceptually consistent, e.g., if both alleles have the same number of CAG repeats, the difference in the transcription rates across alleles has no impact on the overall protein abundance, which is non-trivial for nonlinear models.

The analysis of the estimation results reveals that the three new models with additional mechanisms progressively deepen the suppression of the wt HTT level in the wt/Q111 setting (Fig. 7B-D). Such suppression is achieved when the wt allele HTT level in the wt/Q111 setting is less than half of the level in the wt/wt setting (the region below the half line is thus called the sublinear region). The threshold is at half of the HTT level in wt/wt because wt/Q111 has one instead of two wt alleles. The *Extended Model* is unable to reproduce such suppression, and in fact, the wt allele in the wt/Q111 setting has a higher HTT level. In the model with only nonlinear translation, the simulated wt HTT level in the wt/Q111 setting is still a bit higher than in the wt/wt setting and lies slightly above the half line, outside of the sublinear region. The model with clustering alone produces a downward shift, but combining both non-linearity and clustering yields the strongest suppression effect among the three models, leading the wt HTT level well inside the sublinear region.

**Figure 7.**
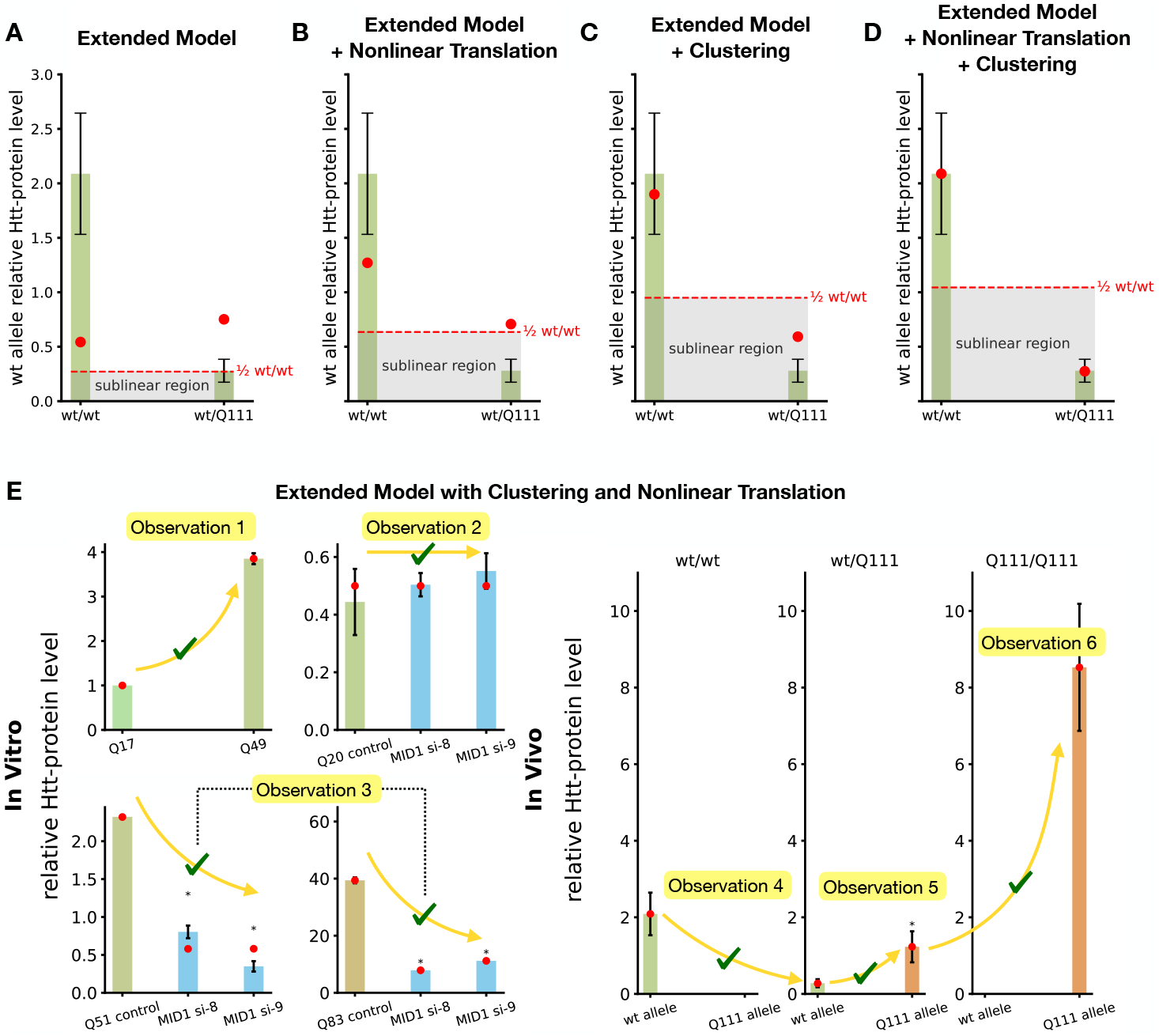
Comparison of model predictions for wild-type allele–derived HTT levels in *wt/wt* and *wt/Q111* settings and fitting results of the *Extended Model with Clustering and Nonlinear Translation*. The HTT levels of the wt allele are indicated for (A) the *Extended Model*, (B) the *Extended Model with Nonlinear Translation*, (C) the *Extended Model with Clustering*, (D) the *Extended Model with Clustering and Nonlinear Translation*. Compared to *Extended Model*, the other three models all improve the fitting regarding the difference in wt allele HTT level between in wt/wt mice and wt/Q111 mice, but only the models with clustering reproduce the sublinear suppression. The red dashed line represents half of the wt allele HTT level of the simulation in the wt/wt setting, and the gray area represents the sublinear HTT level region that is below the half. (E) The *Extended Model with Clustering and Nonlinear Translation* successfully reproduces all observations. The error bars represent the standard deviation of the data. An asterisk above a bar indicates a significant difference (*p <* 0.05) relative to the first bar in the subplot.

Looking at full fitting results of all three models, only the models with clustering can capture all observations (Fig. 7E), while the model with only nonlinear translation does not fully reproduce Observation 4 (see Supplementary Information Sec. S6). Although *Extended Model with Clustering* also captures all observations, only the simulation results of *Extended Model with Clustering and Nonlinear Translation* are within the standard deviation of the data (Fig. 7E).

To quantitatively compare the performance of all models we consider in this work, we calculate the Negative Log-likelihood (NLLH), the Akaike Information Criterion (AIC), its small-sample corrected version (AICc), and the Bayesian Information Criterion (BIC) for each model (Table 1). The *Extended Model with Clustering and Nonlinear Translation* achieves the best fit, as indicated by the lowest NLLH. However, this improved fit requires more free parameters compared to the *Extended Model with Clustering*, which leads to higher complexity penalties and thus worse AIC values. Because our dataset is small (|𝒟| = 48), these penalties are further amplified in AICc and BIC, resulting in the *Baseline Model* scoring best under these criteria. This illustrates an important trade-off: while AICc and BIC naturally prefer simpler models when data is limited, our primary aim here is not parsimony but to identify mechanistic models capable of reproducing all observed data observations. Therefore, the *Extended Model with Clustering and Nonlinear Translation* and the *Extended with Clustering* best fulfill the main purpose of our analysis.

**Table 1:** Overview of the performance of different model topologies. NLLH, AIC, AICc and BIC values are reported – with lower values indicating better performance – as well as the number of model parameters (*n*_*θ*_). Furthermore, it is indicated with a checkmark (✓) if the model reproduces the observations outlined in the text.

| Metric /<br>Observation | Baseline Model | Extended Model | Extended Model + NT | Extended Model + Clustering | Extended Model + Clustering + NT |
| --- | --- | --- | --- | --- | --- |
| **NLLH** | 22.45 | 20.92 | 18.23 | 15.04 | **14.02** |
| **AIC** | 90.90 | 91.83 | 90.47 | **86.08** | 88.04 |
| **AICc** | **136.90** | 150.92 | 166.07 | 171.55 | 197.45 |
| **BIC** | **133.94** | 138.61 | 140.99 | 138.47 | 144.18 |
| $n_\theta$ | 23 | 25 | 27 | 28 | 30 |
| Observation 1 | ✓ | ✓ | ✓ | ✓ | ✓ |
| Observation 2 | ✓ | ✓ | ✓ | ✓ | ✓ |
| Observation 3 | ✓ | ✓ | ✓ | ✓ | ✓ |
| Observation 4 |  |  |  | ✓ | ✓ |
| Observation 5 |  | ✓ | ✓ | ✓ | ✓ |
| Observation 6 | ✓ | ✓ | ✓ | ✓ | ✓ |

**Table 2:**
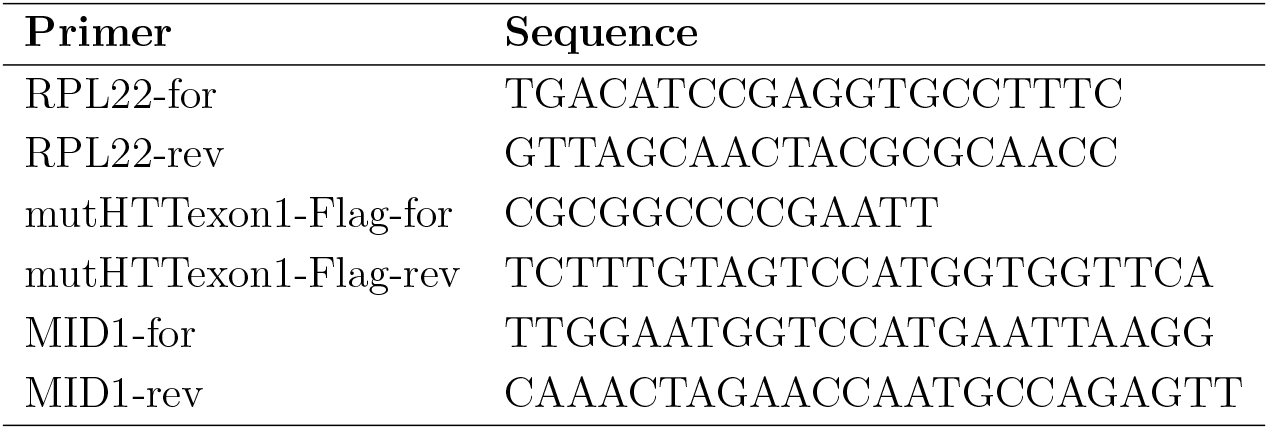
Primer sequences used in this study.

### Simulation study demonstrates how nonlinear translation and clustering help capture all experimentally reported observations

As demonstrated in the last section, the *Extended Model with Clustering and Nonlinear Translation* can reproduce all observations, and its simulation results are within the standard deviation of the data. Specifically, the model can reproduce Observation 4 (the wt allele produces lower HTT levels in wt/Q111 compared to wt/wt mice) and Observation 6 (Q111/Q111 mice show a supralinear increase in HTT levels from the Q111 allele).

To understand how such complex dynamics are captured, we examine how CAG repeat length affects the interactions among the model species and their steady state, demonstrated with the three *in vivo* experiments (Fig. 8A). From wt/wt on the left to Q111/Q111 on the right, more MID1-bound RNA is produced because the binding affinity between MID1 and RNA increases as the CAG repeat length increases. Simultaneously, more MID1-bound RNA can enter the cluster due to the CAG repeat length increase. One can easily see that across three experimental conditions, less and less S6K outside of the cluster gets phosphorylated, with the S6K^P^ level outside being the highest in the wt/wt setting. Less phosphorylation outside of the cluster is due to the increasing binding and clustering with larger CAG repeat length. The increased amount of MID1-bound RNA inside the cluster also means that the level of 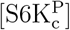 is the highest in the Q111/Q111 setting.

**Figure 8.**
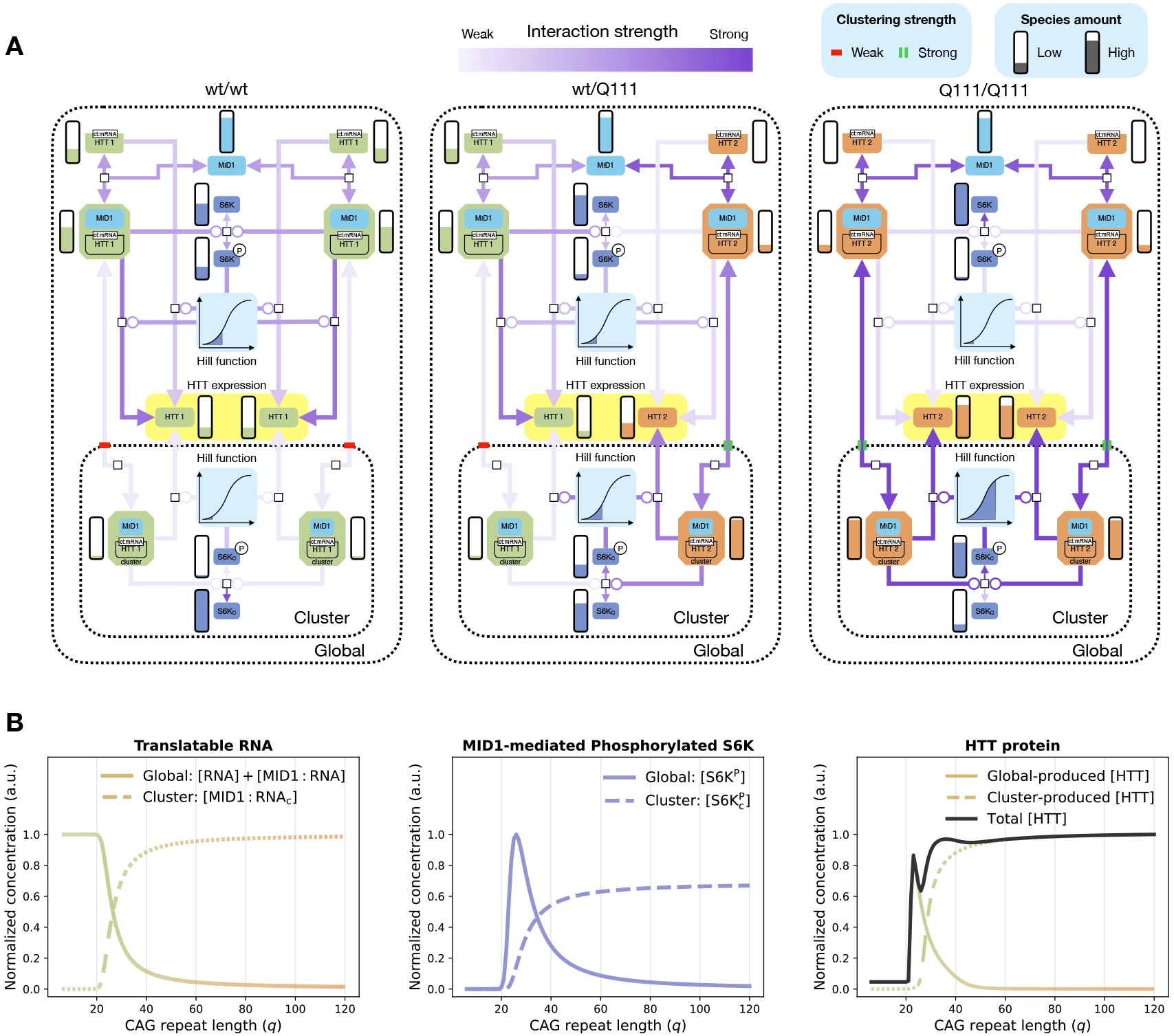
*Extended Model with Clustering and Nonlinear Translation* : simulation study under the three *in vivo* experimental conditions. (A) Comparison of steady state amounts of model species (bars) and interaction strengths in the cluster compartment (purple-gradient arrows) across wt/wt, wt/Q111, and Q111/Q111 settings. CAG repeat length–dependent clustering strength is shown for wild-type (red bar) and mutant (green pass) alleles. Amounts and interactions are from steady state simulations at the maximum likelihood estimate of the model; the Hill function of R3_C,H_ is shown at the cluster center. (B) Steady state simulations for *q*_1_ = *q*_2_ ∈ [6, 120]: left, translatable RNA in global and cluster pools; middle, phosphorylated S6K in both pools; right, HTT levels in each pool and in total. The color gradient represents the CAG repeat length in *Htt* RNA and the Q length in HTT. Curves in each subplot are normalized by the highest value of any quantity in the respective subplot for visual clarity.

The decreasing global S6K^P^ level and increasing cluster S6K^P^ level, together with the nonlinear Hill function, allow *Extended Model with Clustering and Nonlinear Translation* to capture Observation 4. Outside of the cluster, the wt allele in the wt/Q111 setting has a lower translation rate compared to in the wt/wt setting. Meanwhile, although the 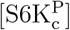 level inside the cluster is higher compared to in the wt/wt setting, the decrease of translation rate in the global environment still dominates. This is because the majority of wt allele translatable resources is in the global environment. Thus, it is possible to reproduce such a suppression phenomenon in the wt/Q111 setting with the drastic decrease in wt allele HTT level outside the cluster and a minor increase inside the cluster.

Similarly, the model can capture the supralinear increase in Q111 allele HTT level in the Q111/Q111 setting compared to the wt/Q111 setting. The minor decrease in translation rate outside the cluster and the fast increase inside the cluster collectively drive the Q111 allele HTT level much higher than in the wt/Q111 setting. Indeed, the spatial segregation yields separate environments, where the nonlinear dependence of translation on S6K^P^ converts the environmental differences into the allele-specific HTT level patterns. CAG repeat length dependent clustering, coupled to a Hill-type translation response, redistributes both molecules and reaction fluxes across the three experimental settings.

To go beyond comparing only the limited number of experimental settings, we systematically explored how different CAG repeat lengths influence the steady state abundance of model species in the *Extended Model with Clustering and Nonlinear Translation*. Specifically, we conduct a simulation study with CAG repeat length between 6 and 120, i.e., *q*_1_ = *q*_2_ and *q*_1_, *q*_2_ ∈ [6, 120]. These simulations reveal that the model predicts that the patterns observed across the three *in vivo* settings (Fig. 8A) are part of a continuous transition, in which translation shifts from being dominated by the global pool at low *q* to being dominated by the cluster pool at high *q* (Fig. 8B, right). Such a shift is caused by two components, translatable RNA and MID1-mediated phosphorylated S6K. Specifically, as CAG repeat length increases, we observe a decrease of global translatable resources and an increase of cluster translatable resources as the resources move into the cluster (Fig. 8B, left). Interestingly, [S6K^P^] does not follow a monotone decrease and has a sharp increase just after *q* = 20 CAG repeats. The sharp increase is due to the shape of the *l* function (see Supplementary Information Sec. S10). Together, those two components drive the HTT curve, global-produced or cluster-produced.

## Discussion

The interaction between RNA and RBP plays a pivotal role in post-transcriptional gene regulation, governing processes such as RNA stability, localization, and translation. Disruptions to these interactions have been increasingly recognized as pathogenic mechanisms, with HD being an example. ^9^ Motivated by experimental data showing a CAG repeat length dependent increase in protein level, we construct a sequence of ODE models to bridge the molecular biochemistry of MID1 binding and S6K phosphorylation with the emergent allele-specific HTT level patterns observed *in vitro* and *in vivo*. The progression—from a minimal *Baseline Model*, through an explicit RNA turnover model (*Extended Model*), to its extensions incorporating nonlinear translation and clustering—allows us to pinpoint which mechanistic ingredients are required to reproduce the full experimental phenotype.

The *Baseline Model*, which encodes only the core known reactions, fails to account for more nuanced observations such as allele-specific HTT level asymmetries. The *Extended Model* extended the *Baseline Model* by adding RNA synthesis and decay. While still incomplete, the model suggests that differences in *Htt* RNA stability, with and without MID1 binding, could explain the preferential translation of the Q111 allele. Indeed, prior work has shown that MID1, besides stimulating translation, may affect RNA stability. ^3^ This prediction is experimentally confirmed in our work: elevated MID1 levels substantially increase the half-life of mutant *Htt* RNA compared to wt. This agreement between model and experiment establishes the *Extended Model* as not only explanatory but also predictive, offering mechanistic insights with direct biological validation.

The prolonged availability of *Htt* RNA due to increased stability through MID1 binding can substantially impact protein synthesis dynamics, potentially exacerbating the accumulation of mutant HTT, a hallmark of HD pathology. Thus, our findings not only reveal a previously unrecognized aspect of RNA regulation by MID1 but also offer valuable insights into the underlying molecular mechanisms contributing to disease progression. Identifying and characterizing this novel MID1-dependent stabilization pathway represents an important step toward developing targeted therapeutic interventions aimed at modulating mutant HTT level through MID1 interaction.

Neither the *Baseline Model* nor the *Extended Model* could explain the suppressed wt HTT level in wt/Q111 (Observation 4), indicating that additional mechanisms may be at play. Firstly, we treat translation downstream of S6K^P^ as cooperative (Hill-type) in the *Extended Model with Nonlinear Translation*. This modification improves the fit to Observation 4 and suggests that S6K^P^ may drive cooperative, threshold-like translational control.

Another extension replaces the well-mixed assumption with a spatially structured description: MID1-bound RNAs can form a local cluster, where S6K phosphorylation and translation are regulated by the local concentrations of MID1:RNA and S6K. This *Extended Model with Clustering* reflects accumulating evidence that many cytoplasmic RNAs reside in sub-cytoplasmic microdomains (e.g., TIS granules/rough endoplasmic reticulum) where translation differs from the surrounding cytosol. ^20^

The concept of a localized cluster with RBP has been studied as a mechanism for protein synthesis.^36^ Our study extends this concept specifically to MID1-mediated clustering of *Htt* RNA. Indeed, MID1 has a known ability to anchor RNA to the microtubule cytoskeleton, facilitating such spatial organization. ^3,2^ Furthermore, as MID1 preferentially binds to RNA with expanded CAG repeats, it is plausible that such clustering is CAG repeat length dependent. This dependence has been shown in other work. ^30^ These findings suggest that subcellular localization and compartmentalized signaling may affect translation efficiency in the context of CAG repeat expansion. As such, disrupting RNA-protein clustering may present a promising therapeutic avenue for modulating allele-specific translation in HD and related disorders.

While the model prediction of MID1-bound RNA clustering is plausible, to validate this mechanistic hypothesis, live-cell imaging and single-molecule RNA visualization would be required, which is beyond the scope of this work. Ideally, the quantitative dependence on CAG-repeat number and MID1 levels should be assessed to recapitulate differences between patients. Such studies would offer critical insights into the spatial architecture of translation regulation in HD.

Finally, the combined *Extended Model with Clustering and Nonlinear Translation* reproduces all observations. To compare the performance of all models, we compute information criteria such as AIC, AICc, and BIC. Although such criteria penalize the additional parameters introduced by the *Extended Model with Clustering and Nonlinear Translation*, the substantially lower NLLH indicates that the model captures observations of the data that simpler models cannot, even if this comes at the cost of increased model complexity. Although the *Baseline Model* has better AICc and BIC measures, the model does not reproduce all observations as well as the *Extended Model with Clustering and Nonlinear Translation*. Given that the purpose of this analysis is to identify mechanistic hypotheses consistent with experimental data observations, we consider that the *Extended Model with Clustering and Nonlinear Translation* still best fulfills the main purpose of our analysis.

In addition to nonlinear translation and clustering, another non-exclusive driver of the morethan-doubling of HTT level in Q111/Q111 could be somatic CAG-repeat expansion. In HD, CAG repeat numbers can increase over time within individual neurons, especially in the striatum, creating mosaic populations with effective repeat lengths well above 111.^16^ Although not explicitly modeled here, for steady state data, this mechanism can be incorporated by estimating a higher-than-111 CAG repeat length.

Although the *Extended Model with Clustering and Nonlinear Translation* captures the key experimental trends, it remains a simplification. We currently represent a single effective cluster, whereas cells may contain multiple, compositionally distinct clusters that could shape translation in parallel. Likewise, we assume one MID1 per RNA, even though expanded *Htt* RNA could recruit multiple MID1 molecules, potentially amplifying clustering and introducing additional cooperativity not captured here.

Beyond these structural assumptions, inference is further limited by data availability: the dataset^27^ is small relative to model complexity, and the wt CAG length in the wt/wt and wt/Q111 cohorts was not reported, requiring inference of *Q*_*wt*_ and adding uncertainty to parameter estimates. This lack of identifiability cautions against favoring one mechanism over another, and we therefore consider all three *Extended Model* with extra components as possible explanations. Future studies with richer time- and dose-response data, explicit reporting of wt repeat lengths, and model extensions (e.g., multiple clusters or multivalent MID1 binding) are needed to improve identifiability and test robustness.

Our integrative modeling and experimental study uncovers a novel mechanistic role for MID1 in promoting both the stability and localized translation of mutant *Htt* RNA through spatial clustering. These findings not only provide insight into how RBP regulates pathogenic RNA behavior in HD but also highlight the critical role of subcellular localization in shaping translation outcomes. By challenging the traditional assumption of a spatially uniform intracellular environment, our work opens new directions for investigating compartmentalized post-transcriptional regulation and lays a foundation for future therapeutic strategies targeting RNA-protein clustering dynamics in neurodegenerative diseases.

## Materials and methods

### Experimental data

We consider previously published experimental data (Fig. 2B). ^27^ A description of the experimental protocols for this published data is in the original publications.^27^

For the new validation experiment we conducted in this study, HEK293 cells stably expressing huntingtin exon1 with 83 CAG repeats in a Tet-off system ^51^ were seeded in poly-L-lysine coated 12 well plates at a density of 2.5 × 10^5^ cells per well. Washing off doxycycline induced the expression of huntingtin exon1. Cells were then cultured for 24 hours and transfected with a plasmid (pCMVTag2A-MID1^2^) to overexpress MID1. After 48 hours of incubation, the actinomycin D time course was started. For this purpose, 1 µl actinomycin D (5 mg/ml; Merck) was added per 1 ml medium, and the cells were harvested after 0 h, 1 h, or 2 h. Cells were washed with PBS and harvested using a cell scraper, followed by total RNA isolation using the Monarch^®^ Total RNA Miniprep Kit (NEB). cDNA was synthesized using the TaqMan reverse transcription reagents kit (Applied Biosystems), and real-time PCR was carried out using the qPCRBIO SyGreen Mix (PCRBiosystems). The sequences of primers are specified in Tab. 2.

### Mathematical formulation

We model the interaction among all the molecules using systems of ODEs. These ODE models are of the form:

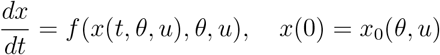

with state vector 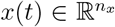 at time *t*, parameter vector 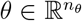, and input vector 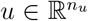 that encodes the experimental conditions. The vector field 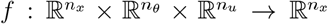 encodes the dynamics of the process and is constructed to be Lipschitz continuous. The function 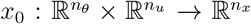 defines the initial condition. By *n*_*x*_, *n*_*θ*_, and *n*_*u*_ we denote the numbers of state variables, unknown mechanistic parameters, and known input variables, respectively.

Model observables are the measurable model quantities. They are defined as functions of the state vector *x* and the parameter vector *θ*

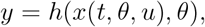

with 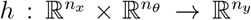. For notational simplicity, we assume that there is only one measurable quantity, i.e., *n*_*y*_ = 1. When the measurements are relative, meaning that they provide data only in a relative form rather than as absolute quantities, it is necessary to introduce scaling factors that scale the observables to the data. For an assumed additive and normally distributed measurement noise, i.e. 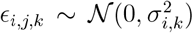, the measurement-toobservable relationship is given by:

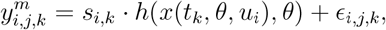

with experimental conditions of each experiment *i* ∈ *{*1, 2, …, *n*_*e*_*}*, repetition index *j* ∈ *{*1 … *n*_*i*_*}*, time point *k* ∈ *{*1, 2, …, *n*_*t*_*}*, and noise standard deviation *σ*_*i,k*_. By *n*_*e*_, *n*_*i*_, and *n*_*t*_ we denote the number of experiments, the number of data replicates and the number of time points. A dataset is defined as the set of all measurements,

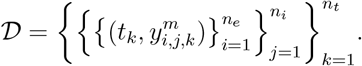

Unknown model parameters (*θ, s, σ*) are inferred from the measured data using maximum likelihood estimation. The likelihood function of observing the experimental data 𝒟 given parameters *θ, s*, and *σ* is given by

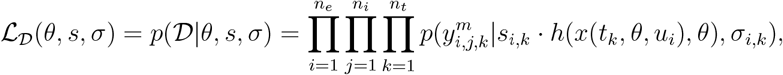

in which the likelihood can be separated into a product of conditional probabilities of individual measurements because of the assumed pairwise independence of all measurements. For a Gaussian noise model, the conditional probability of observing an individual measurement 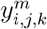 given the observable value of the model *s*_*i,k*_ · *h*(*x*(*t*_*k*_, *θ, u*_*i*_), *θ*) and the noise level *σ*_*i,k*_ is given by

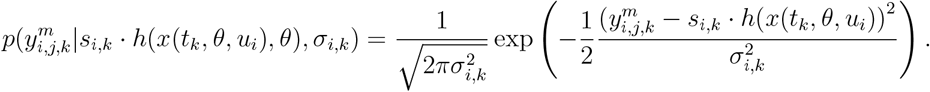

Instead of directly maximizing ℒ_𝒟_, it is equivalent and numerically often preferable to minimize the negative log-likelihood *J* = − log −_𝒟_. Assuming Gaussian noise, this objective function is given by

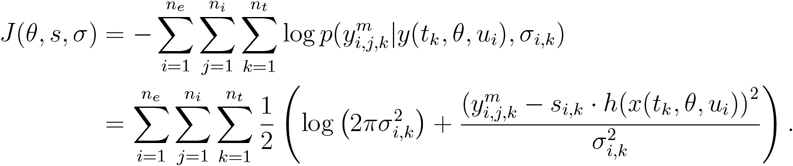

For better convergence and efficiency properties, ^53,28,40^ the unknown model parameters are estimated hierarchically: scaling *s* and noise parameters *σ* of the models are estimated in the inner optimization problem, and all other model parameters *θ* are estimated in the outer optimization problem. The hierarchical optimization problem is given by

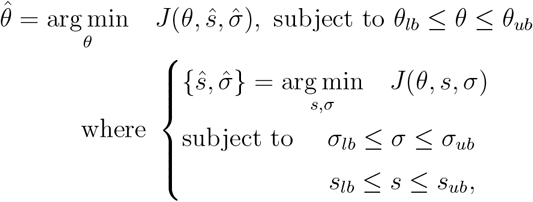

in which the inner optimization problem can be solved analytically. ^40^

The selection of the bounds for the unknown model parameters (*θ, s, σ*) is outlined in the Supplementary Information (see Supplementary Information Sec. S3).

### Steady state computation

A steady state of an ODE model 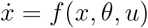 is defined as a state vector *x*^*^ satisfying

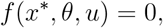

i.e., all time derivatives vanish. Numerically, steady states can be obtained in different ways. One common approach is to directly solve the nonlinear system *f* (*x, θ, u*) = 0 using a Newton-type root-finding method, which requires evaluating and factorizing the Jacobian *∂f/∂x*. Alternatively, the system can be integrated forward in time until the norm of *f* (*x, θ, u*) falls below a predefined tolerance, indicating proximity to a steady state. In practice, simulation tools, including the one used in this study, alternate between these steady state calculation approaches for increased robustness.^12^ In all cases, suitable convergence criteria (e.g., relative and absolute tolerances) must be specified to determine when the system is sufficiently close to steady state.

### Model selection criteria

We employ the Akaike Information Criterion (AIC), ^1^ its small-sample corrected version (AICc),^22^ and the Bayesian Information Criterion (BIC) ^43^ as model selection metrics to assess the trade-off between goodness-of-fit and model complexity. These criteria are defined as:

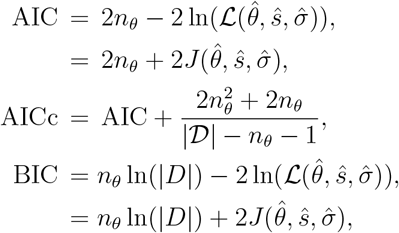

where 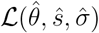 is the maximum likelihood value, and |𝒟| is number of data points.

### Uncertainty quantification

Here, we define what it means for a parameter vector to lie in a confidence region of a certain significance. This is done using the likelihood-ratio test, in which the corresponding test statistic is defined as

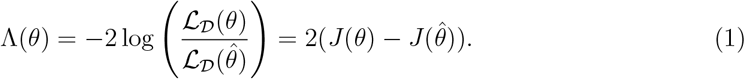

In the asymptotic case of a large number of data points, the test statistic converges to a chi-square distribution 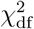 with df = *n*_*θ*_ degrees of freedom (see^50^ for more details). Then the confidence region of significance *α* is defined as

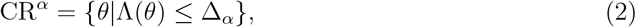

in which Δ_*α*_ is the *α*-th percentile of the 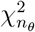 distribution. The one-dimensional projection of a confidence region for a specific parameter *θ*_*m*_ of the parameter vector *θ* is defined as the Confidence Interval (CI) of that parameter:

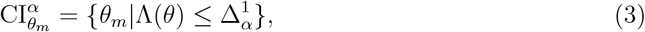

in which 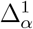 is the *α*-th percentile of the 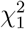 distribution with one degree of freedom. Prediction uncertainty quantification can then be carried out by propagating parameter uncertainty using the CR to model predictions.

## Implementation and availability

We use PEtab^41^ to define the parameter estimation problems and SBML^21^ for standardized representations of the mathematical models. We create the SBML models using Tellurium’s Antimony.^7^ Parameter optimization and uncertainty analysis are implemented using pyPESTO 0.5.6,^39^ wherein we use AMICI 0.33.0^12^ for simulation and steady state calculation, and Fides 0.7.8^13^ for optimization. Each model is estimated using gradient-based multi-start local optimization with 50000 randomly sampled starting points. As is commonly done, most parameters are estimated on the logarithmic scale to uniformly explore parameter orders of magnitude.^17^ All SBML and PEtab files, and the code and instructions to perform all analyses, are available at https://zenodo.org/records/17478762.

## Author Contributions

J.H., S.K., and T.T. conceptualized the project and acquired financial support. Y.L., J.H., and T.T. developed the mathematical model. Y.L., D.D., and J.H. calibrated and analyzed the model. S.K. and A.R. conducted the wet-lab experiments. J.H., Y.L., and D.D. wrote the initial draft and prepared the visualization. All authors discussed the results and commented on the manuscript.

## Supplementary information

### S1 ODE systems and initial conditions

Here we present in detail the dynamical system, the initial condition, and the parameter vector of all models.

#### S1.1 Baseline Model

The *Baseline Model* considers the four core reactions (R1-R4) mentioned in the main manuscript, which involve the following state vector:

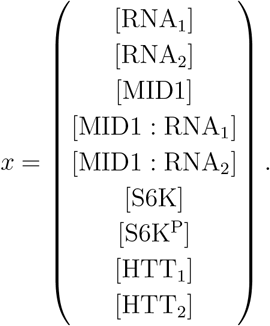

We assume the initial condition, i.e., *x*_0_, is as follows:

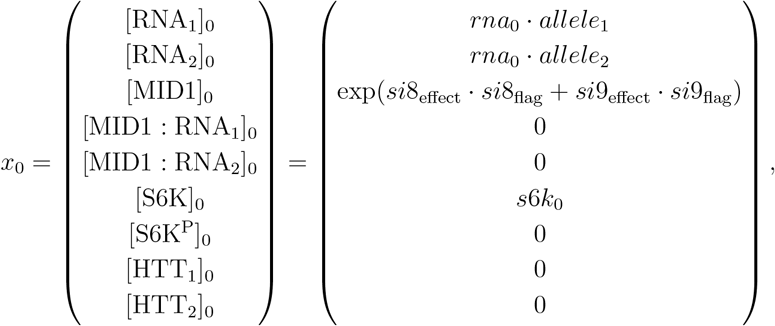

Here, *rna*_0_, *s*6*k*_0_, *si*8_effect_ and *si*9_effect_ are parameters to be estimated, while *allele*_1_, *allele*_2_, *si*8_flag_ and *si*9_flag_ are part of the experimental condition (see Sec. S2). In the *Baseline Model*, we assume the initial RNA level is a constant, contrary to the *Extended Model* (see Sec. S1.2). The binary values of *allele*_1_ and *allele*_2_ indicate whether the experiment is performed *in vitro* (e.g., *allele*_1_ = 1 and *allele*_2_ = 0) or *in vivo* (*allele*_1_ = *allele*_2_ = 1). In some experiments, MID1 is knocked down to assess its effect on the HTT level using small interfering RNA (siRNA) techniques-specifically with si8 or si9. The flags *si*8_flag_ and *si*9_flag_ indicate when a specific type of knockdown is applied, and the corresponding effects are captured by the parameters *si*8_effect_ and *si*9_effect_. Because the intracellular concentration of S6K is unknown, the initial conditions *s*6*k*_0_ is also treated as an estimated parameter.

Given the reactions (R1-R4), we can write down the ODE for each species according to the reactions in which it participates. Combining these equations yields the complete ODE system, ∀*t* ∈ [0, ∞)

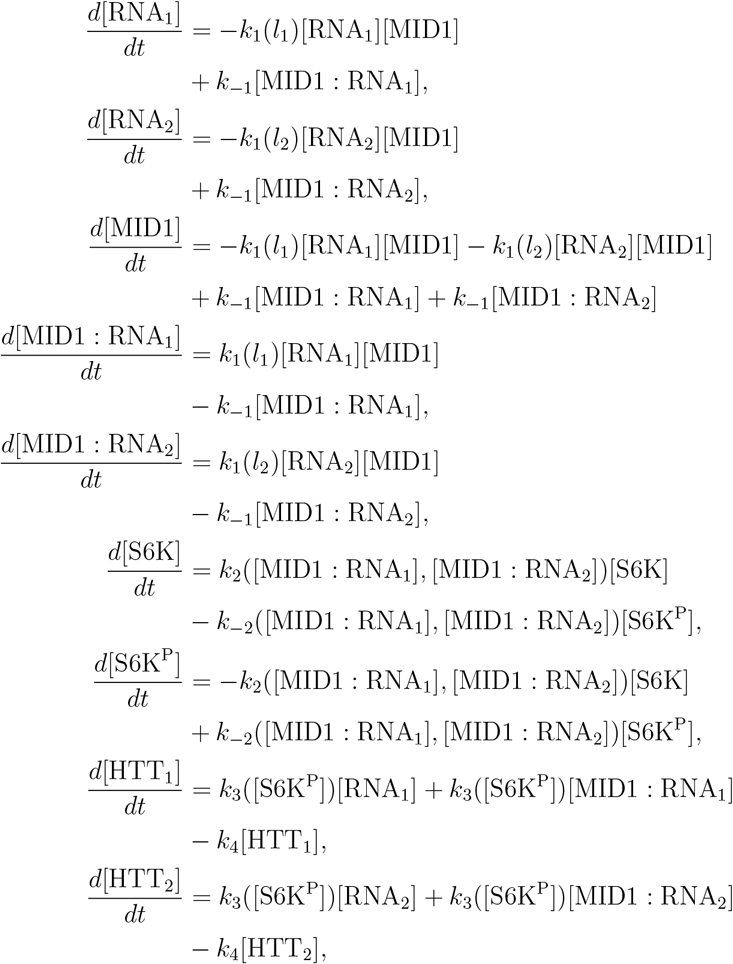

where,

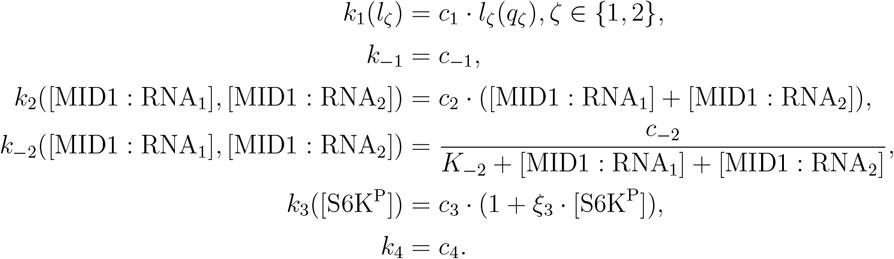

Based on the above ODE system, we point out that the system is symmetric with respect to RNA_1_ and RNA_2_, meaning switching the two variables would not change the model dynamics.

We combine all parameters used in this work in one table (see Sec. S3), and the parameters that are part of the *Baseline Model* can be written in the vector form:

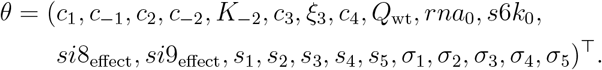

The scaling parameters (*s*_1_, …, *s*_5_) are introduced in the main text and discussed in more detail in Sec. S3. The details on noise parameters (*σ*_1_, …, *σ*_5_) are included in Sec. S11.

#### S1.2 Extended Model

Since the additional three reactions (R5-R7) do not add additional species, the state vector of *Extended Model* stays the same as in the *Baseline Model*. However, the initial condition *x*_0_ is now changed to:

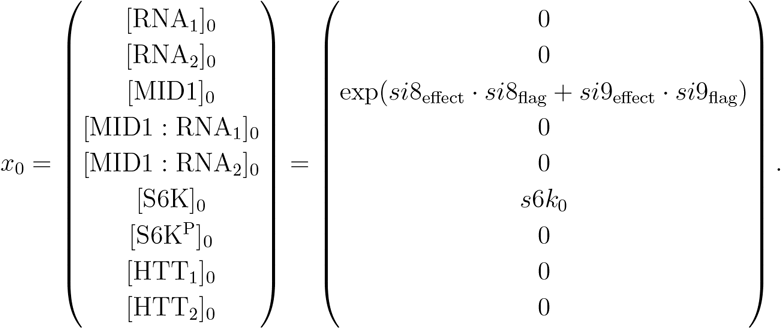

In the *Baseline Model*, we assume the same RNA initial conditions, [RNA_*ζ*_]_0_ = *s*6*k*_0_, across all experimental conditions. Because there is no RNA degradation reaction and [MID1 : RNA_*ζ*_]_0_ = 0, the [RNA_*ζ*_]+[MID1 : RNA_*ζ*_] should be conserved to *rna*_0_ under all experimental conditions and the four reactions. In the *Extended Model*, such conservation does not exist any more because two reactions, RNA production and RNA degradation, are added. For simplicity, we assume the initial condition of RNA is 0.

Given the reactions (R1-R7), the complete ODE system is as follows, ∀*t* ∈ [0, ∞):

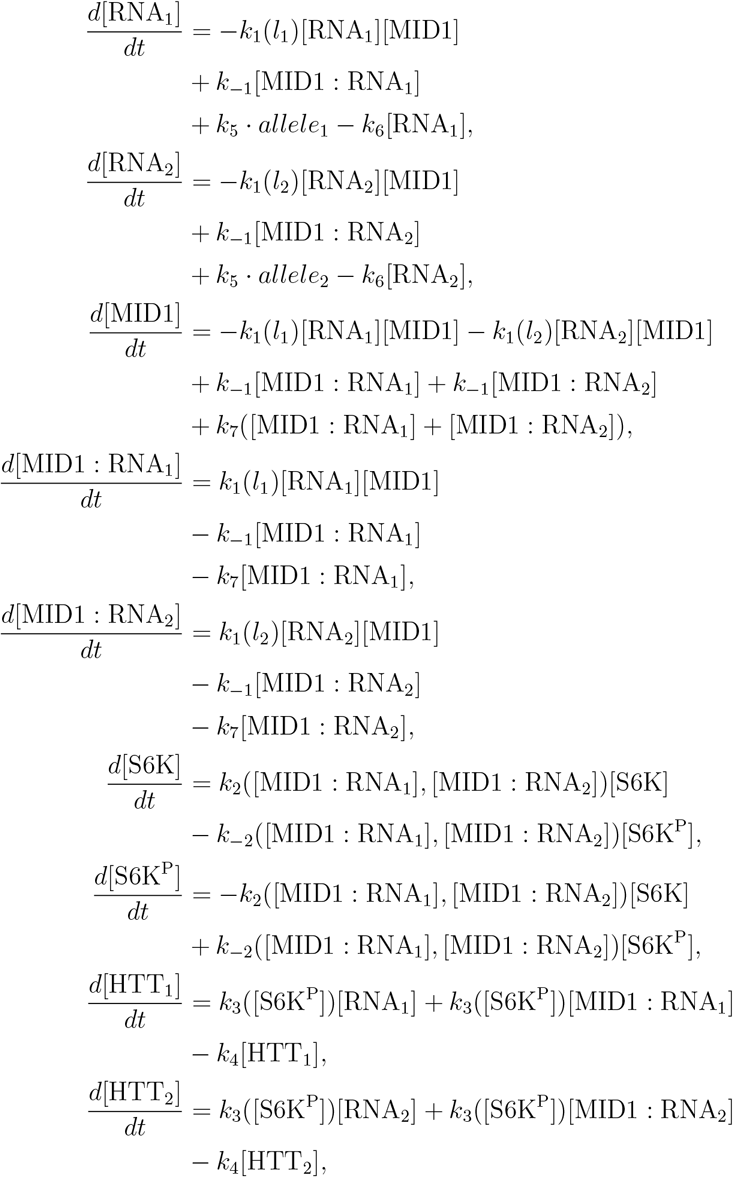

where,

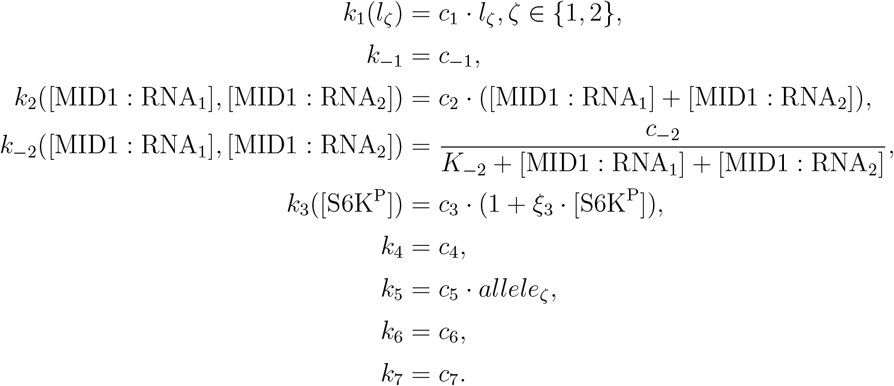

Notice that the experimental condition indicating *in vitro* or *in vivo* experiment, *allele*_*ζ*_, is now part of the transcription rate, *k*_5_ · *allele*_*ζ*_. The control with *allele*_*ζ*_ on whether one allele produces RNA or not would also subsequently control whether other reactions of that allele happen or not.

Because of the extra three reactions, three parameters are added and *rna*_0_ is removed. The new parameter vector is:

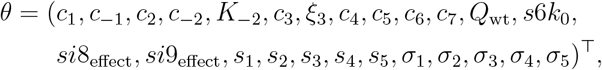

#### S1.3 Extended Model with Nonlinear Translation Rate

The state vector, initial condition, and ODE systems of the *Extended Model with Nonlinear Translation* are the same as in the *Extended Model*, with only *k*_3_ changed to *k*_3,*H*_ (R3_H_):

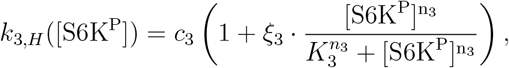

and *k*_6_ changed to *k*_6,*R*_ = *ξ*_6_ · *k*_7_, where 1 ≤ *ξ*_6_ (R6_R_). Due to the change in these two functions, the parameter vector is updated to include *n*_3_, *K*_3_, *ξ*_6_:

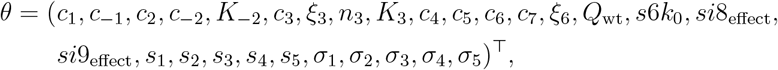

#### S1.4 Extended Model with Clustering (*l*-dependent)

The *Extended Model with Clustering* (*l*-dependent) shares the same reactions (R1-R7) with the *Extended Model*, with one reaction modified (R6_R_) and four new reactions (R8, R2_C_, R3_C_, R7_C_) inside the cluster. The additional reactions introduce four additional states inside the cluster. The state vector is as follows:

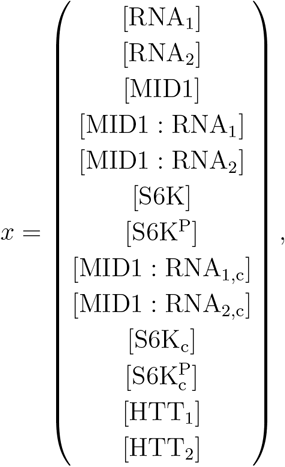

with initial condition *x*_0_ being:

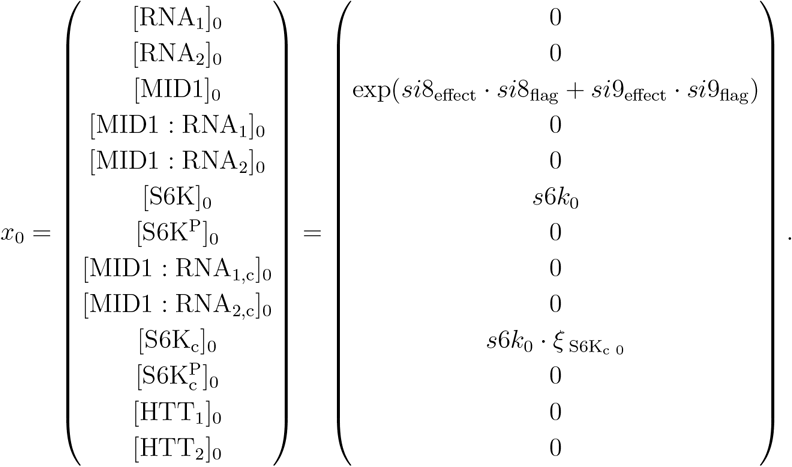

Notice the extra parameter 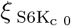. Because the cluster forms as more MID1:RNA is being produced, the concentration of S6K inside and outside the cluster can be dynamic. However, as we only have steady-state measurements, we assume the conserved pool of S6K and S6K_c_ are constant. We define the constant ratio between [S6K_c_]_0_ and [S6K]_0_ as 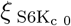.

Given all the reactions, the complete ODE system is as follows, ∀*t* ∈ [0, ∞):

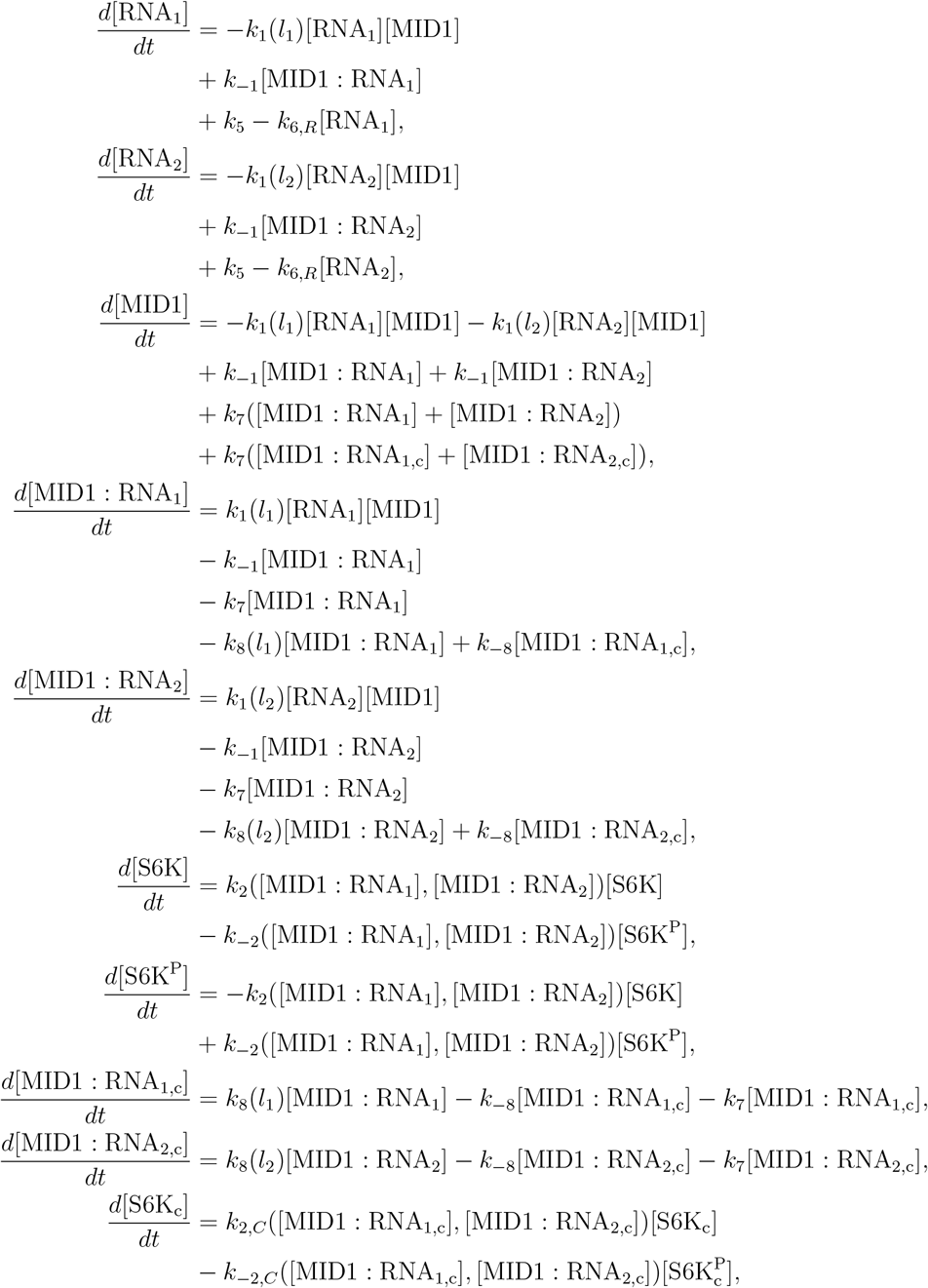

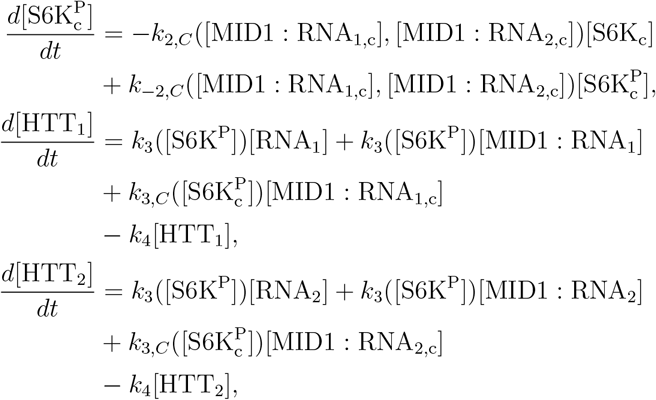

where,

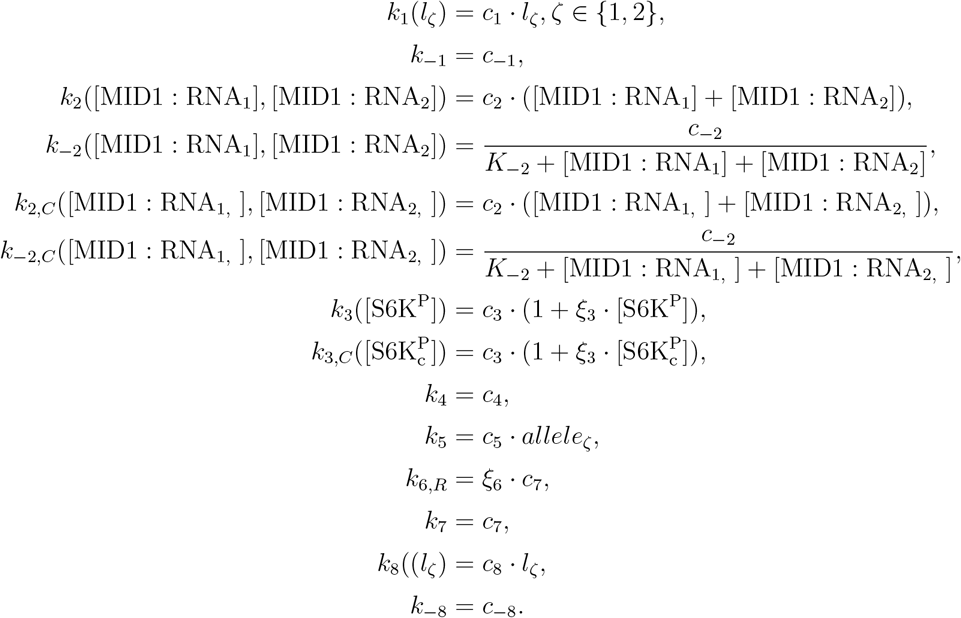

Notice that *k*_2_ and *k*_2,*C*_ (same for *k*_−2_ and *k*_−2,*C*_ or *k*_3_ and *k*_3,*C*_) share the same parameters, but the inputs of the two functions are different.

Due to the extra reactions, the parameter vector is now as follows:

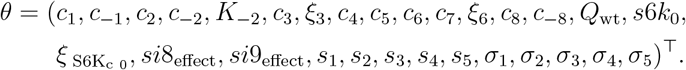

#### S1.5 Extended Model with Clustering and Nonlinear Translation (*l*-dependent)

The *Extended Model with Clustering and Nonlinear Translation* (*l*-dependent) shares the same state vector, initial condition, and ODE system of the *Extended Model with Clustering (l-dependent)*, with only the *k*_3_ changed to *k*_3,*H*_ outside the cluster (R3_H_):

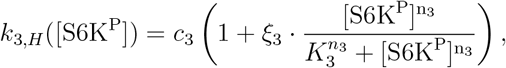

and *k*_3,*C*_ changed to *k*_3,*C,H*_ (R3_C,H_):

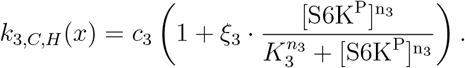

The corresponding parameter vector with the extra two parameters, *n*_3_ and *K*_3_, is:

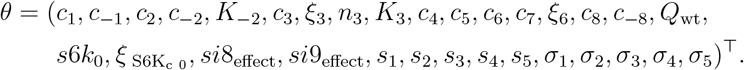

#### S1.6 Extended Model with Clustering (non-*l*-dependent)

In the main manuscript, we only discuss in detail the models with clustering where the clustering rate is *l* dependent, *Extended Model with Clustering* and *Extended Model with Clustering and Nonlinear Translation*. To show the effect of *l* dependence, we also explore the models without clustering, where the rate is not *l* dependent-*Extended Model with Clustering* (non-*l*-dependent) and *Extended Model with Clustering and Nonlinear Translation* (non-*l*-dependent). We compare the four models to demonstrate why *l* dependence might be necessary to explain all data observations in Sec. S6.1.

The *Extended Model with Clustering* (non-*l*-dependent) shares the same state vector, initial condition, ODE system, and the parameter vector of the *Extended Model with Clustering (l-dependent)*, with only *k*_8_ changed to *k*_8_(*l*) = *c*_8_.

#### S1.7 Extended Model with Clustering and Nonlinear Translation Rate (non-*l*-dependent)

The *Extended Model with Clustering and Nonlinear Translation* (non-*l*-dependent) shares the same state vector, initial condition, ODE system, and parameter vector of the *Extended Model with Clustering and Nonlinear Translation* (*l*-dependent), with only *k*_8_ changed to *k*_8_(*l*) = *c*_8_.

### S2 Experimental Condition

The dataset^1^ comprises 14 experiments, of which 11 are *in vitro* and 3 are *in vivo*. Each experiment is characterized by six binary or numerical variables, which we denote as experimental conditions 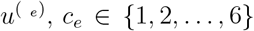 and collect into a vector as input 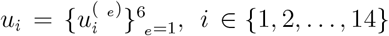. The six experimental conditions are: (1) *u*^(1)^ = *allele*_1_: whether allele 1 expresses (binary), (2) *u*^(2)^ = *allele*_2_: whether allele 2 expresses (binary), (3) *u*^(3)^ = *q*_1_: the CAG repeat length of allele 1 (non-negative), (4) *u*^(4)^ = *q*_2_: the CAG repeat length of allele 2 (non-negative), (5) *u*^(5)^ = *si*8_flag_: whether siRNA si-8 is applied (binary), (6) *u*^(6)^ = *si*9_flag_: whether siRNA si-9 is applied (binary). Note that if *u*^(1)^ = 0 (*u*^(2)^ = 0), then *u*^(3)^ = 0 (*u*^(4)^ = 0) as the corresponding allele is not expressed.

The experimental conditions influence the dynamics in three main ways: allele expression, CAG repeat length, and MID1 knockdown. First, if an allele is expressed, transcription for that allele is “on,” with the rate implemented as *k*_5_ = *c*_5_ · *u*^(*ζ*)^ = *c*_5_ · *allele*_*ζ*_. In the *Baseline Model*, where RNA levels are constant, the initial RNA level is set to [RNA]_0_ = *rna*_0_ · *u*^(*ζ*)^ = *rna*_0_ · *allele*_*ζ*_. For a non-expressing allele, this value is zero. Second, the values *q*_1_ and *q*_2_ (*u*^(3)^ and *u*^(4)^) enter the system through the *l* function (see Sec. S4) Third, the Application of siRNA si-8 or si-9 reduces MID1 levels. This is modeled by setting the initial condition for MID1 as [MID1]_0_ = exp(*si*8_effect_ · *u*^(5)^ + *si*9_effect_ · *u*^(6)^) = exp(*si*8_effect_ · *si*8_flag_ + *si*9_effect_ · *si*9_flag_). The specific values of these six experimental conditions for each experiment are listed in Tab. S1.

### S3 Complete Parameter Table

We summarize all parameters used in each model in our work in the Tab. S2. In addition to the kinetic parameters, initial conditions, and noise parameters, our framework includes five independent scaling factors (*s*_1_, …, *s*_5_), each corresponding to a distinct western plot gel on which subsets of experiments were measured. Experiments run on the same gel share a scaling factor, as their band intensities are directly comparable, while experiments from different gels require separate scaling factors to account for inter-gel variation. For example, Q17 and Q49 originate from the same gel and thus share one scaling factor, whereas Q20, Q20 si8, and Q20 si9 were measured together on another gel and share a different scaling factor.

**Table S1:**
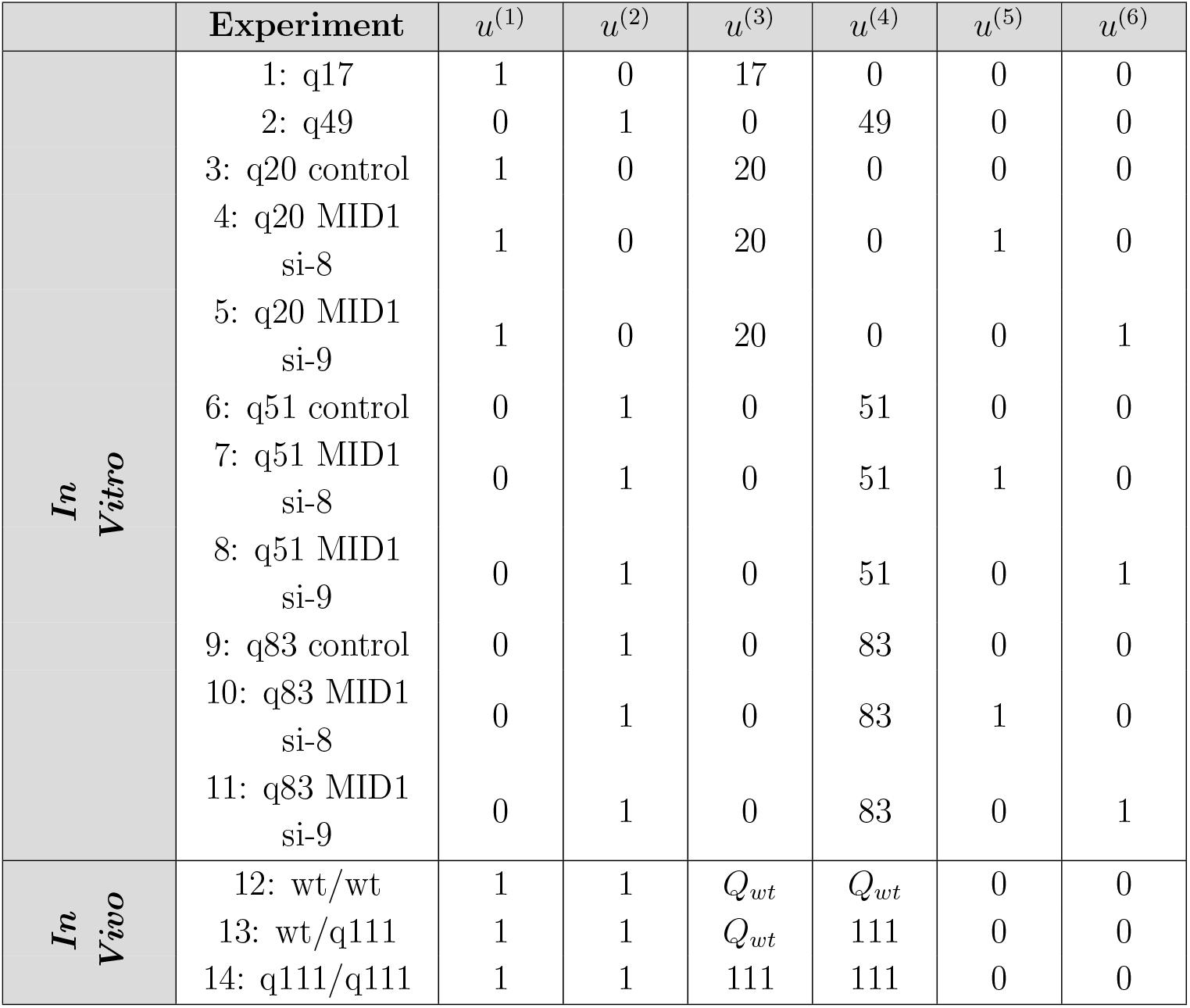
Experimental conditions. Experimental configurations of data, ^1^ showing experimental conditions 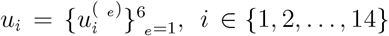 in each experiment.

### S4 Activation function, *l*, **as a function of CAG repeat length** *q*

To model how CAG repeat length (*q*) affects the binding affinity function *k*_1_ and clustering rate function *k*_8_, we use hairpin length (Fig. S1C), constructed to be a function of *q*, as part of *k*_1_ and *k*_8_ functions. We use hairpin length, rather than *q* directly, because MID1 binds with RNA through the stable hairpin structure. Without the hairpin structure, which can happen when *q* is very small, the binding affinity with MID1 and thus the clustering rate might be very weak or non-existent.

Based on the in silico RNA-structure study of CAG repeat stretches with different repeat sizes,^1^ hairpin length changes very little when *q* is small, while the hairpin length linearly increases after a certain *q* threshold (Fig. S1A). A linear function of *q* can not reproduce such dynamics. We thus model hairpin length in the form of an “activation function”:

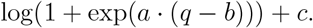

**Table S2:** Parameters. Parameter, their associated reaction or definition, bounds, and prior distribution used for optimization. All parameters are fitted. *Units:* UC = unit of concentration; UT = unit of time; “–” = dimensionless.

| Reaction or Definition | Name | Lower Bound | Upper Bound | Prior | Unit |
| --- | --- | --- | --- | --- | --- |
| Binding | $c_1$ | 1E–6 | 1E8 | log-uniform | $UC^{-1} \cdot UT^{-1}$ |
| Unbinding | $c_{-1}$ | 1E–6 | 1E8 | log-uniform | $UT^{-1}$ |
| Phosphorylation | $c_2$ | 1E–6 | 1E8 | log-uniform | $UC^{-1} \cdot UT^{-1}$ |
| Unphosphorylation | $c_{-2}$ | 1E–6 | 1E8 | log-uniform | $UC \cdot UT^{-1}$ |
| | $K_{-2}$ | 1E–14 | 1E8 | log-uniform | UC |
| Translation | $c_3$ | 1E–6 | 1E8 | log-uniform | $UT^{-1}$ |
| | $\xi_3$ | 1E0 | 1E2 | log-uniform | – |
| | $n_3$ | 1 | 4 | uniform | – |
| | $K_3$ | 1E–6 | 1E8 | log-uniform | UC |
| HTT Degradation | $c_4$ | 1E–6 | 1E8 | log-uniform | $UT^{-1}$ |
| Transcription | $c_5$ | 1E–6 | 1E8 | log-uniform | $UC \cdot UT^{-1}$ |
| RNA Degradation | $c_6$ | 1E–6 | 1E8 | log-uniform | $UT^{-1}$ |
| | $\xi_6$ | 1E0 | 1E5 | log-uniform | – |
| MID1:RNA Degradation | $c_7$ | 1E–6 | 1E8 | log-uniform | $UT^{-1}$ |
| Clustering | $c_8$ | 1E–6 | 1E8 | log-uniform | $UT^{-1}$ |
| Declustering | $c_{-8}$ | 1E–6 | 1E8 | log-uniform | $UT^{-1}$ |
| wt CAG repeat length | $Q_{wt}$ | 7 | 36 | uniform | – |
| [RNA]<sub>0</sub> | $rna_0$ | 1E–6 | 1E8 | log-uniform | UC |
| [S6K]<sub>0</sub> | $s6k_0$ | 1E–6 | 1E8 | log-uniform | UC |
| [S6K<sub>c</sub>]<sub>0</sub>/[S6K]<sub>0</sub> | $\xi_{[S6K_c]_0}$ | 1E–3 | 1E0 | log-uniform | – |
| siRNA si-8 effect | $si8_{effect}$ | –10 | 0 | uniform | – |
| siRNA si-9 effect | $si9_{effect}$ | –10 | 0 | uniform | – |
| scaling factor 1 | $s_1$ | 1E–6 | 1E8 | uniform | – |
| scaling factor 2 | $s_2$ | 1E–6 | 1E8 | uniform | – |
| scaling factor 3 | $s_3$ | 1E–6 | 1E8 | uniform | – |
| scaling factor 4 | $s_4$ | 1E–6 | 1E8 | uniform | – |
| scaling factor 5 | $s_5$ | 1E–6 | 1E8 | uniform | – |
| noise parameter | $\sigma_1$ | 0 | ∞ | uniform | – |
| noise parameter | $\sigma_2$ | 0 | ∞ | uniform | – |
| noise parameter | $\sigma_3$ | 0 | ∞ | uniform | – |
| noise parameter | $\sigma_4$ | 0 | ∞ | uniform | – |
| noise parameter | $\sigma_5$ | 0 | ∞ | uniform | – |

**Figure S1:**
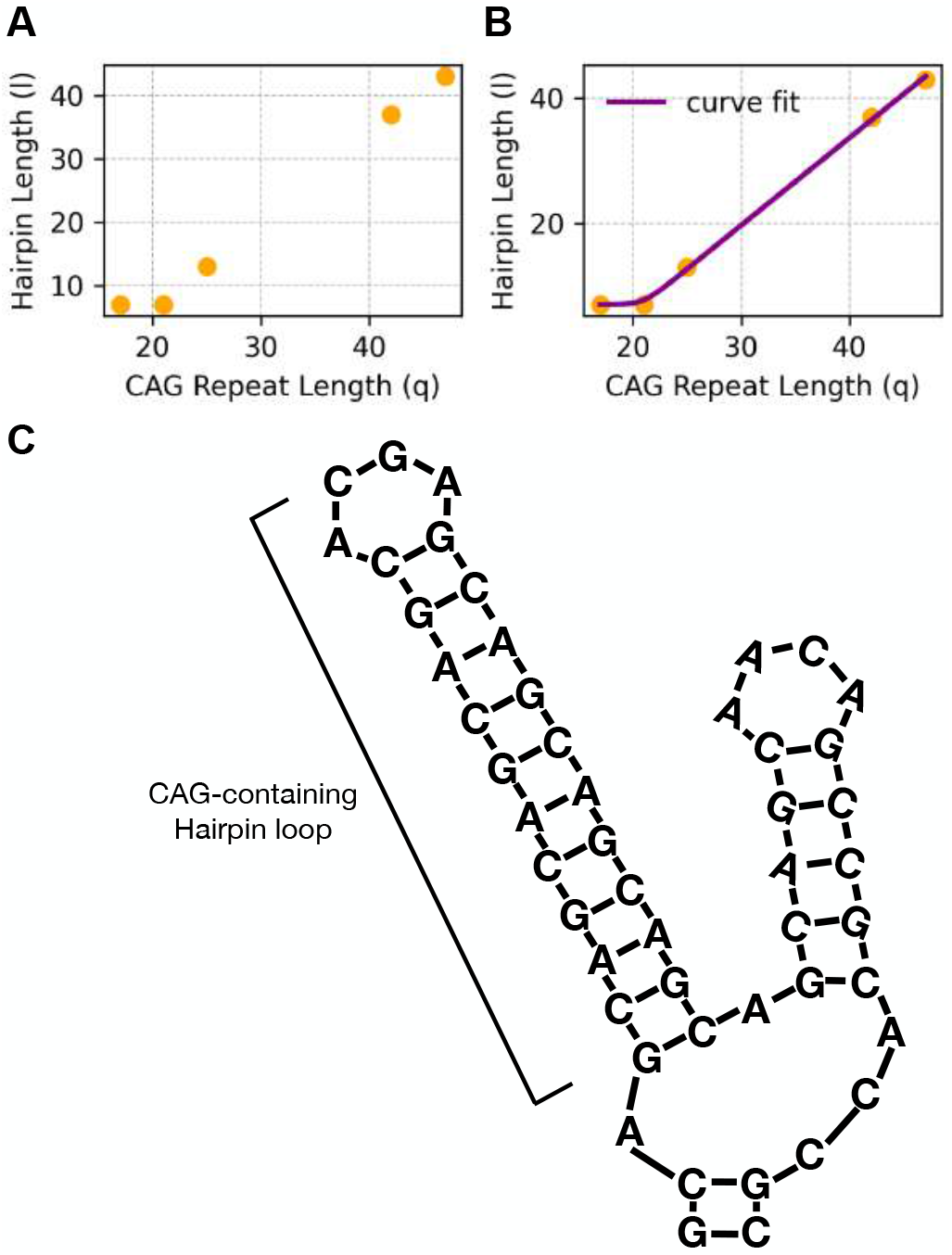
Relationship between CAG repeat length and hairpin structure length. (A) Hairpin length (*l*(*q*)) plotted against CAG repeat length (*q*), based on an in silico RNA secondary-structure prediction study. ^1^ The hairpin length shows minimal change at low CAG repeat numbers but increases linearly beyond a threshold repeat length. (B) Curve fitting with an “activation function” to capture the nonlinear relationship between CAG repeat length and hairpin length. (C) Representative illustration of a CAG-containing RNA hairpin structure, highlighting the formation of stable secondary structures responsible for MID1 binding affinity and RNA clustering.

However, notice that the hairpin length is seven for both 17 CAG and 21 CAG repeat length. To reduce the number of parameters, we let *c* = 7. Choosing (*a* = 1.45, *b* = 30) to be the initial guess of the optimization, the curve fitting result (Fig. S1B) is:

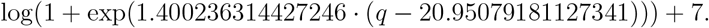

As the binding affinity or the clustering rate should possibly be non-existent when *q* is small, we offset the hairpin length function and define *l* to be:

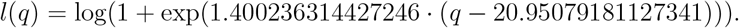

The binding affinity and clustering rate are thus in the form of:

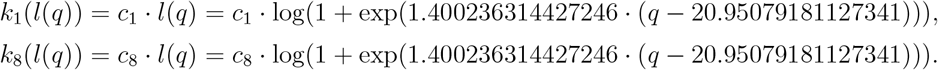

## S5 Existence, uniqueness, and non-negative invariance of ODE solutions proof

### Proposition.

*Consider an ODE system*

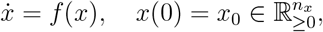

*with the following properties:*

### (A1)Regularity

*f is C*^1^ *on an open set containing* 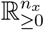

### (A2)Inward boundary

*For each i, f*_*i*_(*x*) ≥ 0 *whenever x*_*i*_ = 0 *and* 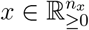

### (A3)Uniform bounds

*There exist nonnegative linear functionals T*_*k*_(*x*) = *α*_*k*_ · *x and constants a*_*k*_ ≥ 0, *b*_*k*_ *>* 0 *such that*

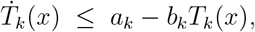

*and the T*_*k*_ *collectively bound all coordinates*.

*Then the initial value problem admits a unique global solution* 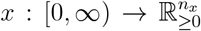, *and the nonnegative orthant is forward invariant*.

*Sketch of proof*. By (A1), *f* is locally Lipschitz, so there is a unique local solution (Picard– Lindelöf). By (A2), the vector field points inward at each boundary face, hence solutions starting nonnegative remain nonnegative. By (A3), Grönwall’s inequality implies uniform bounds on each *T*_*k*_, and thus on all components of *x*(*t*). The trajectory remains in a compact, positively invariant set on which *f* is globally Lipschitz. Therefore, local solutions extend uniquely for all *t* ≥ 0.

#### Remark.

*In the models in this work, assumptions (A1)–(A3) are satisfied as follows:*

- *Regularity (A1) holds since all right-hand sides consist of polynomials or smooth rational/Hill-type functions with strictly positive denominators*.
- *Inward boundary condition (A2) is guaranteed because all degradation or loss terms are proportional to the state variable itself and vanish at the boundary, while formation terms remain non-negative*.
- *Uniform bounds (A3) are obtained from biologically meaningful totals (e*.*g. conserved pools of S6K, MID1, or RNA) and clearance terms that yield differential inequalities of the form* 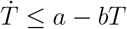.

*Therefore, each specific model variant is a special case of the proposition*.

## S6 Comparison of fitting results of all models

To examine which model performs the best, we check the fitting results (Fig. S2), which are usually the most important indicator. Visual inspection of the fits reveals that the model incorporating both clustering and nonlinear translation—particularly the *Extended Model with Clustering and Nonlinear Translation* (*l*-dependent)—captures the observed patterns most accurately across all experimental conditions. In contrast, simpler models (e.g., the *Baseline Model*) misestimate HTT level in several conditions.

Model selection results using NLLH, AIC, AICc, and BIC (Tab. S3) provide a quantitative perspective on model performance. While the *Extended Model with Clustering and Nonlinear Translation* (*l*-dependent) achieves the lowest negative log-likelihood (NLLH = 14.02), indicating the best likelihood-based fit, the *Extended Model with Clustering* (*l*-dependent) without nonlinear translation achieves the best score in AIC. Other criteria, AICc and BIC, prefer simpler models, such as the *Baseline Model*. This discrepancy arises because NLLH reflects pure fit quality without penalizing complexity, while AIC, AICc and BIC severely penalize the additional parameters introduced by nonlinear translation and clustering.

### S6.1 *l* dependence is necessary for clustering rate

Interestingly, models without *l*-dependent clustering fail to or only barely capture the sublinear decrease in HTT level in wt/Q111 mice (Fig. S3). This failure to fit Observation 4 may indicate that CAG repeat length–dependent clustering is a necessary feature to reproduce the full spectrum of observed behaviors. Indeed, this differential clustering efficiency is shown to be critical in *Extended Model with Clustering and Nonlinear Translation* (*l*-dependent). As wt MID1-bound RNA is effectively excluded from the cluster compared to Q111 MID1-bound RNA in wt/Q111 mice, such RNA cannot collaboratively enhance the local S6K phosphorylation that occurs within the cluster, constraining the translation rate inside the cluster to not increase too much. Meanwhile, wt HTT level becomes sublinear in wt/Q111 setting due to wt MID1-bound RNA’ restricted localization to the global compartment, where translation rate is at a lower basal rate. This basal translation level in wt/Q111 mice is lower compared to that observed in wt/wt mice, primarily because Q111 MID1-bound RNA is largely located inside the cluster.

**Figure S2:**
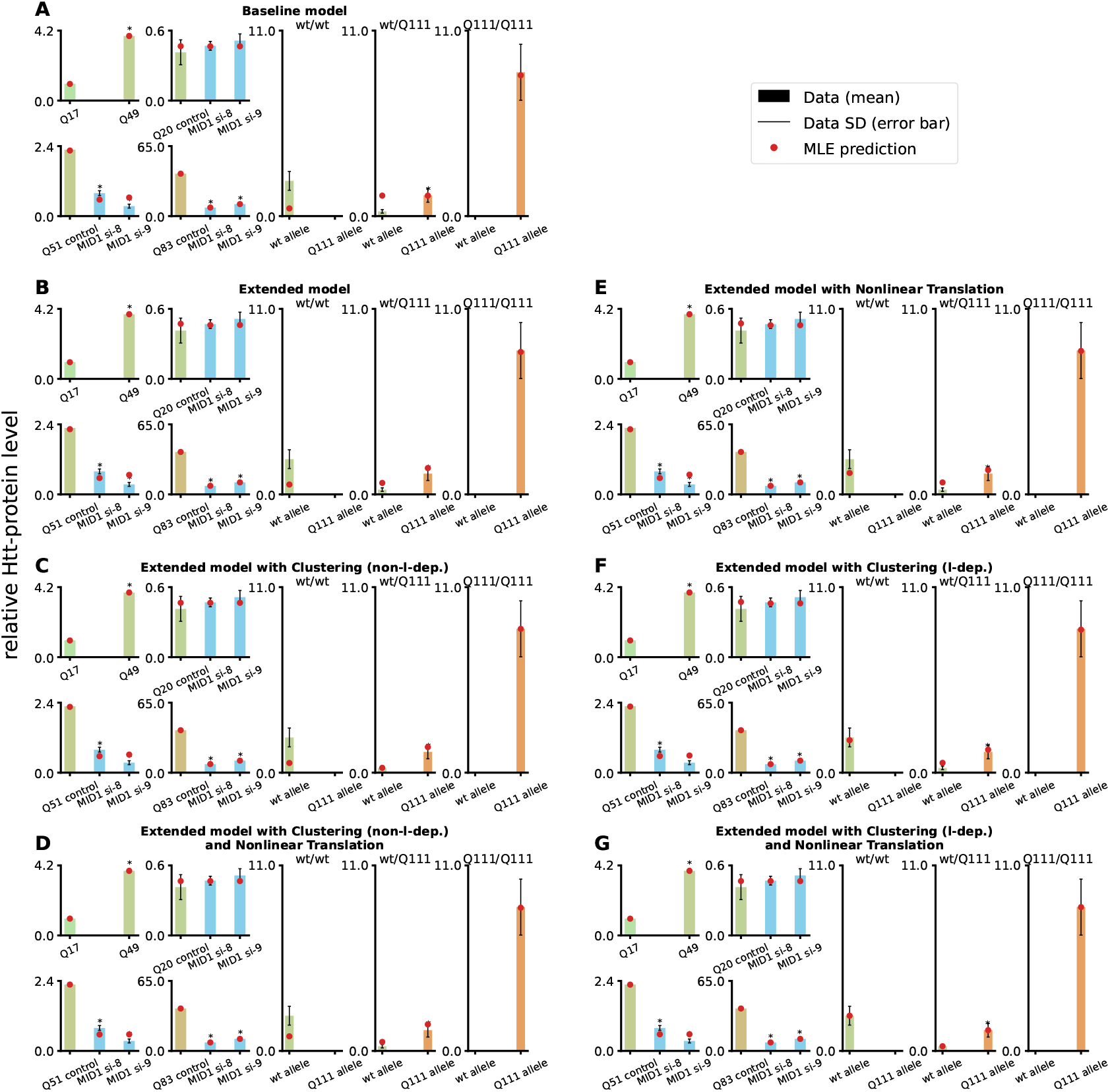
Model fits for all models. Bars indicate the mean relative HTT level (black line: standard deviation), with model predictions at maximum likelihood estimates (red dots). Panels compare the *Baseline Model, Extended Model, Extended Model with Nonlinear Translation, Extended Model with Clustering, Extended Model with Clustering and Nonlinear Translation*, and variants incorporating clustering without CAG repeat length dependence (non-*l*-dep.). Models with *l*-dependent clustering show improved agreement with the data, particularly in wt/wt and wt/Q111 settings.

**Table S3:** Model comparison of all models using NLLH, AIC, AICc, and BIC. Lower values indicate better performance. *n*_*θ*_ indicates the number of parameters. NT indicates Nonlinear Translation.

| Model | NLLH | AIC | AICc | BIC | $n_\theta$ |
| --- | --- | --- | --- | --- | --- |
| Baseline Model | 22.45 | 90.90 | **136.90** | **133.94** | 23 |
| Extended Model | 20.92 | 91.83 | 150.92 | 138.61 | 25 |
| Extended + NT | 18.23 | 90.47 | 166.07 | 140.99 | 27 |
| Extended + Clustering (non-$l$-dep.) | 19.60 | 95.20 | 180.68 | 147.60 | 28 |
| Extended + Clustering ($l$-dep.) | 15.04 | **86.08** | 171.55 | 138.47 | 28 |
| Extended + Clustering + NT (non-$l$-dep.) | 17.95 | 95.91 | 205.32 | 152.04 | 30 |
| Extended + Clustering + NT ($l$-dep.) | **14.02** | 88.04 | 197.45 | 144.18 | 30 |

The fact that the *l*-dependent clustering models compared to their respective variants without *l*-dependent clustering rank higher in both likelihood and penalized criteria highlights the importance of this mechanism in explaining the data. Models lacking *l*-dependence or clustering generally perform worse in all metrics, further reinforcing that length-dependent clustering is a key mechanistic feature.

Recent experimental work has reported CAG repeat length–dependent homotypic RNA clustering,^2^ a phenomenon conceptually consistent with the clustering mechanism explored in our models. While those findings were observed within condensates, our framework suggests that analogous repeat-length–dependent clustering could arise in localized cytoplasmic translation hubs, where MID1-bound RNA may influence both spatial organization and local S6K^P^–dependent translation. Such parallels offer an intriguing indication that length-dependent clustering could be a relevant regulatory feature in the context of Huntington’s disease, although further experimental validation will be required.

## S7 Steady-state RNA and MID1:RNA level comparison between wt and Q111 alleles

We examined the steady-state total RNA levels (sum of free and MID1-bound species) for both alleles in the *Extended Model*. As shown in Fig. S4, the Q111 allele consistently reaches a higher translatable resource abundance than the wt allele under identical transcription rates. This result provides a mechanistic explanation for the competitive expression pattern observed *in vivo*: the lower degradation rate of MID1-bound RNA (*c*_7_) compared to free RNA (*c*_6_) leads to preferential stabilization of Q111 RNA, which binds MID1 more strongly. The temporal simulation (left panel) shows this difference emerging over time, while the bar plot (right panel) summarizes the steady-state values, highlighting the persistent imbalance.

**Figure S3:**
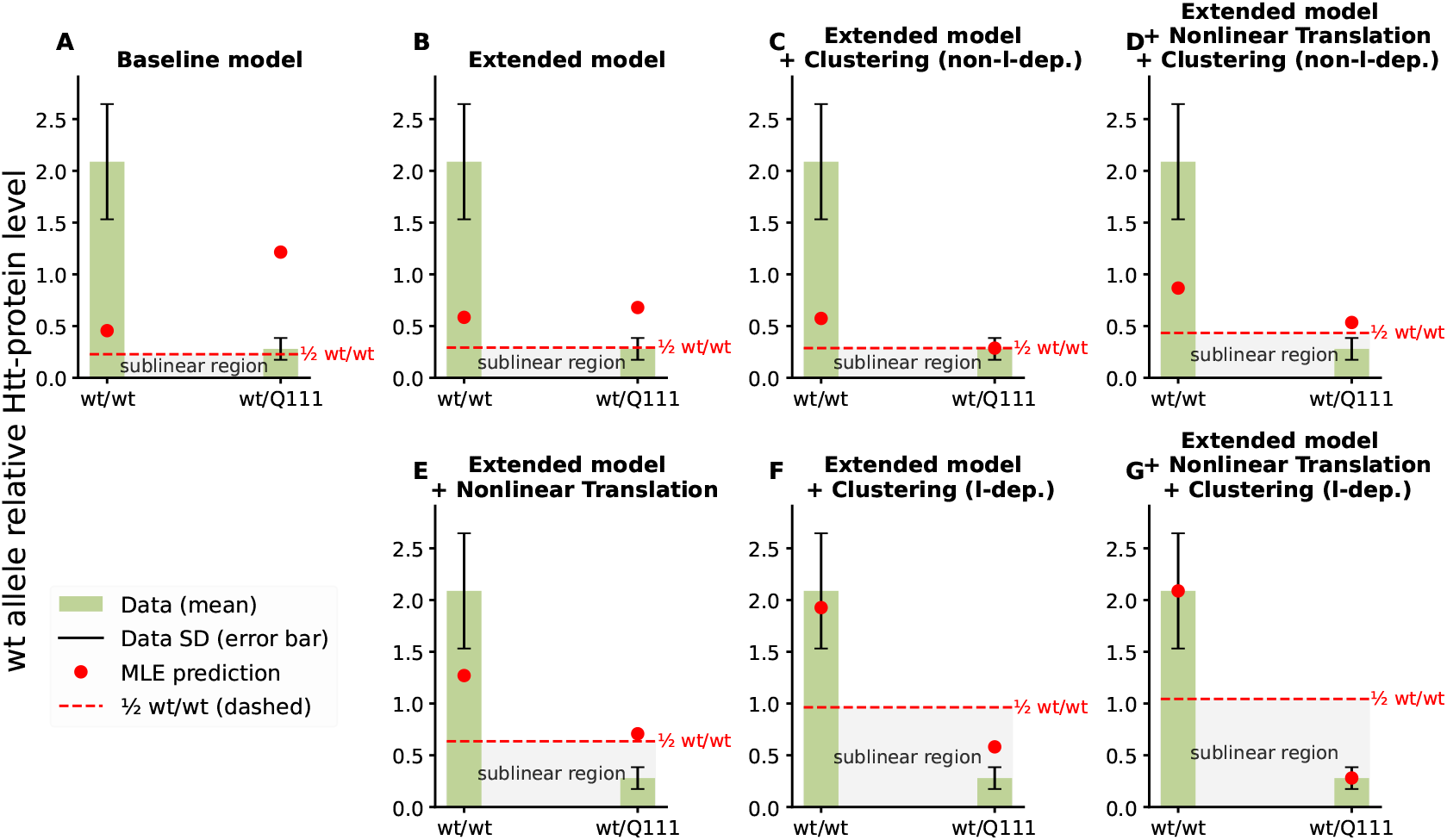
Comparison of model predictions for wild-type allele–derived HTT levels in *wt/wt* and *wt/Q111* settings. Green bars indicate experimental mean values with standard deviation (black error bars), red dots represent model predictions at maximum likelihood estimates, and the dashed red line marks half the predicted *wt/wt* HTT level. The shaded “sublinear region” highlights HTT levels below the half threshold. Panels show *Baseline Model, Extended Model, Extended Model with Nonlinear Translation, Extended Model with Clustering, Extended Model with Clustering and Nonlinear Translation*, and variants incorporating clustering without CAG repeat length dependence (non-*l*-dep.). Models with *l*-dependent clustering better reproduce the sublinear HTT level in *wt/Q111* mice.

**Figure S4:**
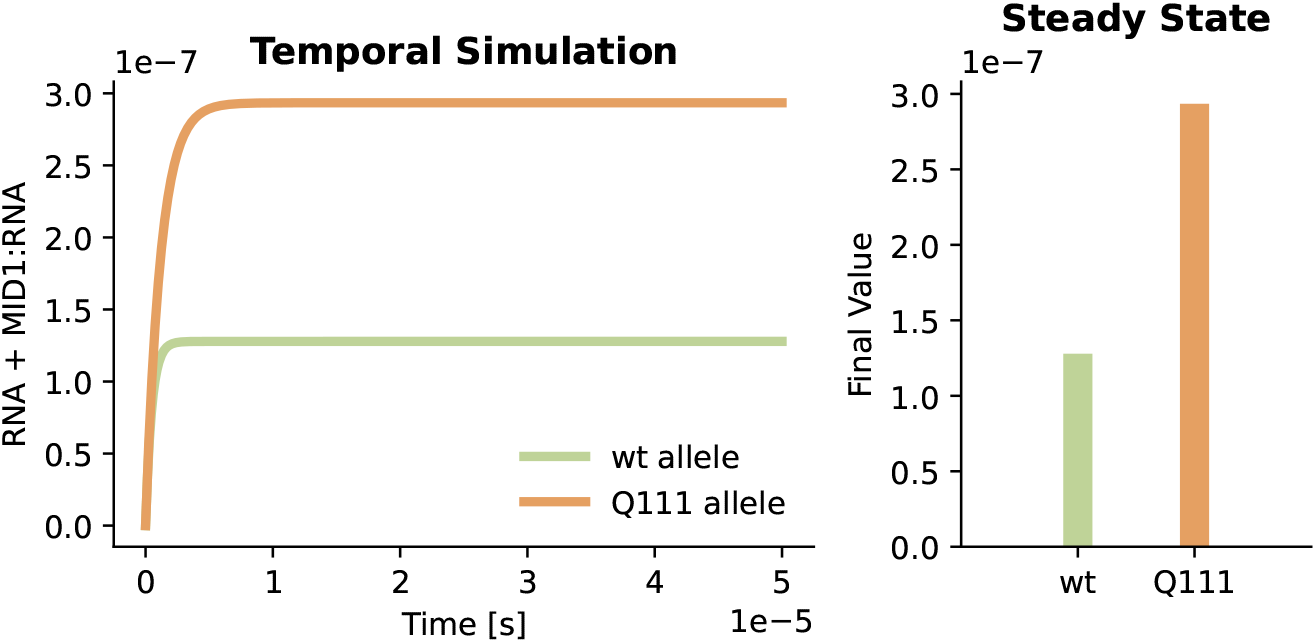
Comparison of total RNA + MID1:RNA levels for wt and Q111 alleles in the *Extended Model*. Left: Time-course simulation showing total RNA (free and MID1-bound) for wt (green) and Q111 (orange) alleles under identical transcription rates. Right: Corresponding steady-state values, indicating consistently higher RNA abundance for Q111 allele. This imbalance arises from the lower degradation rate of MID1-bound RNA (*c*_7_) relative to free RNA (*c*_6_), leading to preferential stabilization of Q111 RNA.

## S8 Relationship between *k*_6_ and *k*_7_ in *Extended Model* s with extra components

In the *Extended Model*, parameter estimates consistently indicated that the degradation rate of the MID1:RNA complex (*k*_7_) was lower than the degradation rate of RNA alone (*k*_6_), suggesting a stabilizing effect of MID1 on RNA. This prediction was experimentally confirmed. Consequently, for all subsequent models, we constrained this relationship by setting *k*_6_ = *ξ*_6_ · *c*_7_ with *ξ*_6_ *>* 1.

To investigate how this constraint is manifested in different model structures, we examined the distribution of (*k*_6_, *k*_7_) pairs seen across all optimization histories for the five extended models. For each model, we retained only points within the 95% confidence region and plotted them alongside the maximum likelihood estimate (MLE) (Fig. S5).

All models retain a shape of confidence regions similar to that observed for the *Extended Model*, characterized by an elongated ridge parallel to the identity line. This indicates an identifiable ratio of *k*_6_ and *k*_7_, but a nonidentifiability of the individual parameters. Furthermore, for *Extended Model with Nonlinear Translation* the MLE lies away from the identity line, as it does for the *Extended Model* (Fig. S5 A, D). In contrast, all clustering models (Fig. S5 B, C, E, F) show MLEs located closer to the identity line, implying that clustering tends to prefer smaller differences between *k*_6_ and *k*_7_.

**Figure S5:**
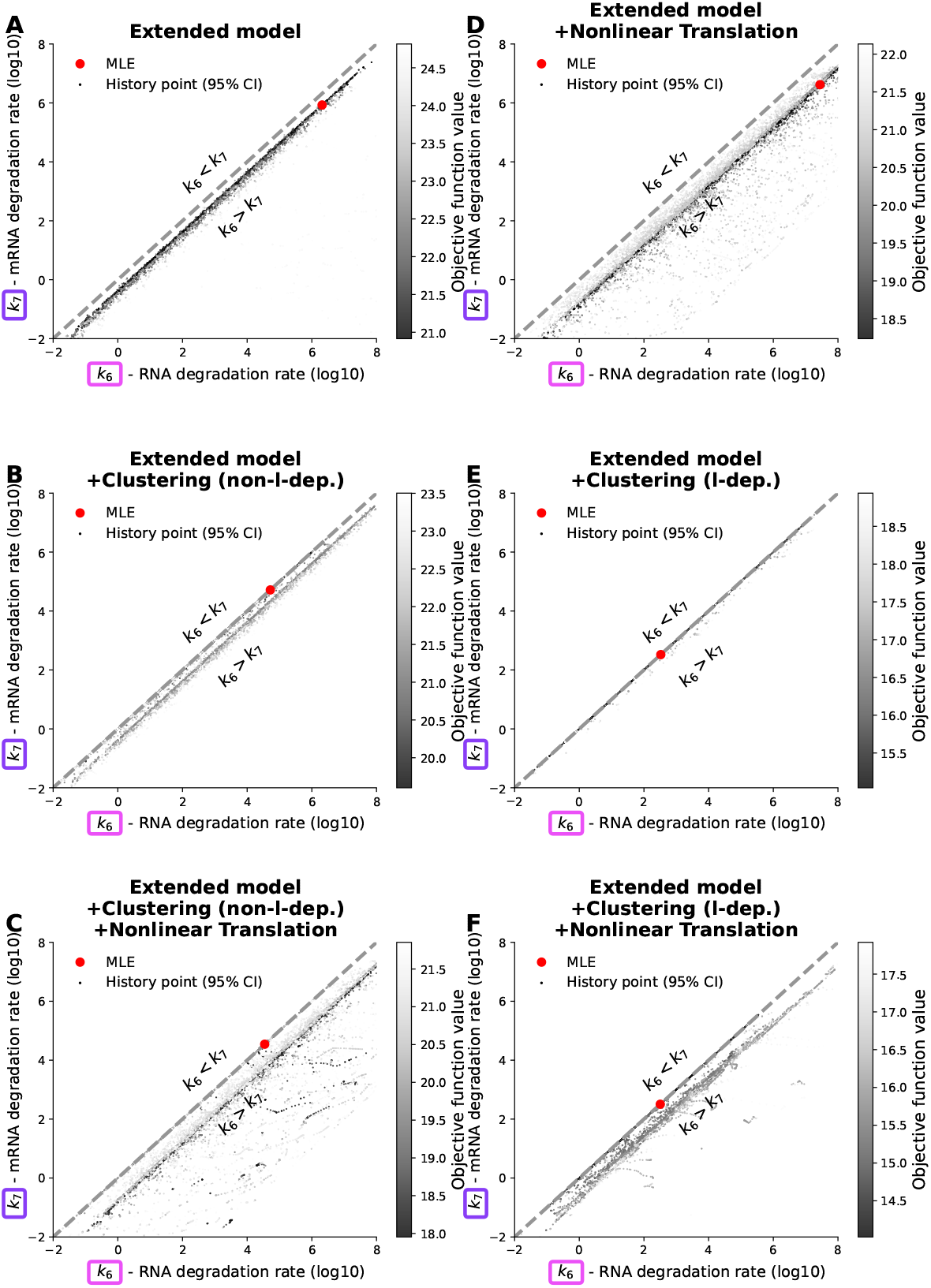
Scatter plots of (*k*_6_, *k*_7_) reaction rate pairs for all models. Parameters (log10 scale) are obtained from optimization history for the *Extended Model, Extended Model with Nonlinear Translation, Extended Model with Clustering*, and *Extended Model with Clustering and Nonlinear Translation*, and variants incorporating clustering without CAG repeat length dependence (non-l-dep.). The maximum likelihood estimates are shown in red. Darker points indicate better fits within the 95% confidence region.

However, even in models where the MLE lies close to the identity line, there remain parameter vectors within the 95% confidence region where *ξ*_6_ *>* 2 (equivalently, log *ξ*_6_ *>* 0.4), reflecting a high difference between *k*_6_ and *k*_7_. The objective function values for these points are very similar to the MLE, with ΔNLLH *<* 0.3.

## S9 Optimized *Q*_*wt*_ distribution

In the experimental conditions wt/wt and wt/Q111, the CAG repeat length of the wild-type allele is not experimentally determined. We therefore treat this quantity, denoted *Q*_*wt*_, as a free parameter to be estimated during model calibration. Fig. S6 summarizes the maximum likelihood estimate (MLE) and 95% confidence intervals (CI) of *Q*_*wt*_ for all model variants. CIs were obtained using parameter values of the endpoints of model optimization that had a final value of the objective function below the CI threshold.

For simplicity in estimation, we treat *Q*_*wt*_ as a continuous rather than a discrete quantity, even though CAG repeat numbers are inherently integer valued. This choice has a negligible impact on model behavior, as the hairpin length function *l*(*q*) used in all model variants is continuous in *q*, ensuring that small fractional differences in *Q*_*wt*_ produce smooth and well-defined effects on downstream dynamics.

The 95% confidence intervals indicate a high uncertainty of the parameter estimate for most models, some even covering the whole estimation space. However, the uncertainty is reduced for the two models with the best objective function values – the *Extended Model with Clustering (l-dep*.*) and Nonlinear Translation* and the *Extended Model with Clustering (l-dep*.). An interesting thing to point out is that the MLE, the best parameter value obtained, for these models is just above 20 repeats, from ~21 to ~23.

## S10 Effect of the Hill function on translation dynamics across CAG repeat lengths

In the simulation study of the *Extended Model with Clustering and Nonlinear Translation* (l-dependent) (Fig. S7), as the CAG repeat length increases, the total HTT level follows an overall increasing trend, but there are local minima at which the HTT level is lower, despite a higher CAG repeat length. This dynamic is driven by the opposing trends of global and cluster MID1-mediated S6K^P^ levels: as *q* increases, [S6K^P^] declines and 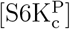 increases, with the Hill-type translation dependence converting these changes into compartment-specific translation rates.

To understand the HTT level dynamics described above, we examined the Hill function for both the global and cluster compartments at selected CAG repeat lengths: low translation in both compartments (*q* = 20), peak translation in the global compartment (*q* = 23), peak translation after first drop (*q* = 36) and second local minimum of translation (*q* = 46) (Fig. S8).

**Figure S6:**
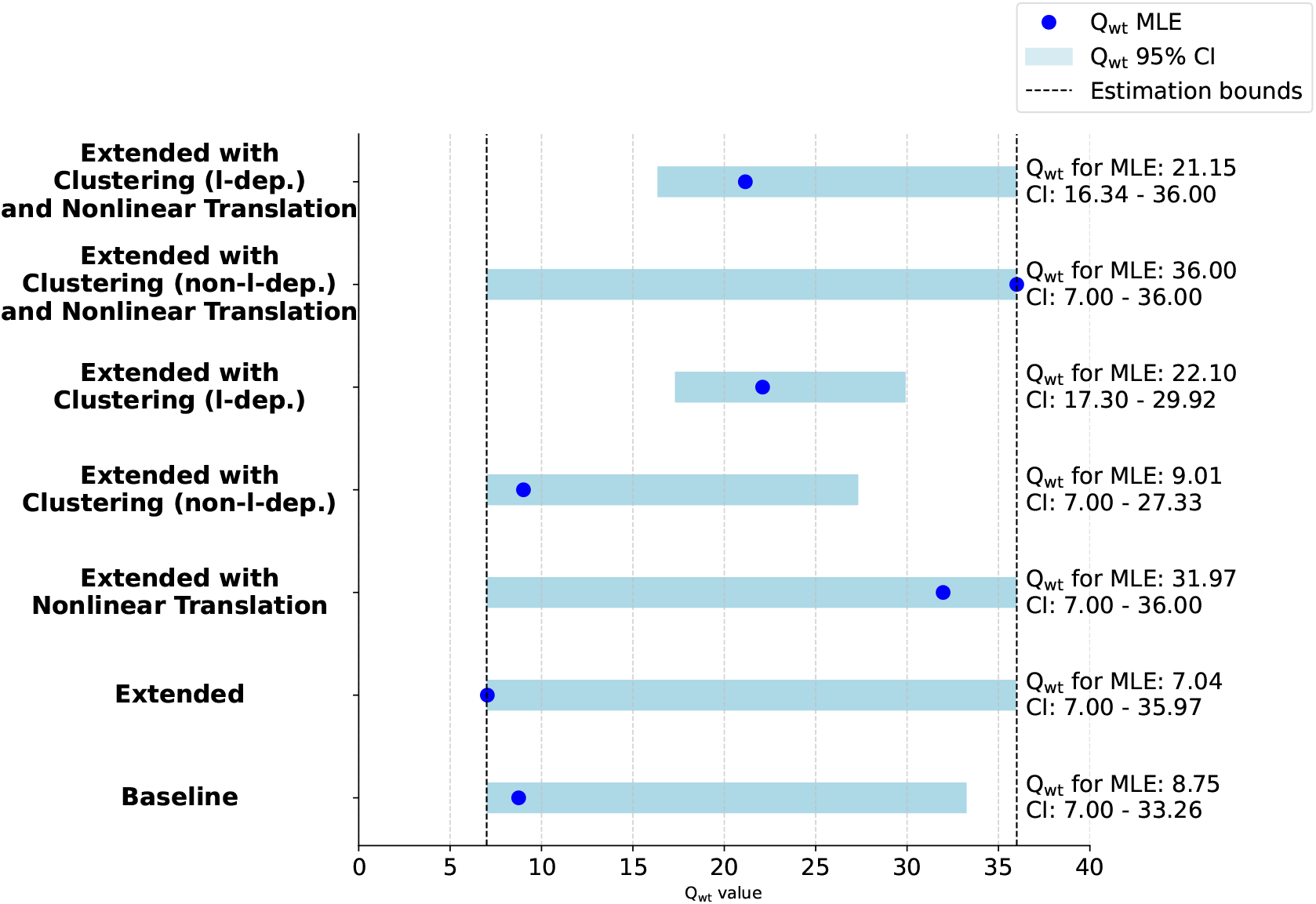
Estimated *Q*_*wt*_ across models. Blue dots indicate the maximum likelihood estimate (MLE) of *Q*_*wt*_; black dashed lines mark the estimation bounds. Black error bars represent 95% confidence intervals, calculated from optimization endpoint distributions.

Before *q* = 20, the threshold of the *l* function causes the plateau regions for all species, i.e., the steady state of all species does not change when CAG repeats increase. At *q* = 20, there is minimal binding of MID1 to RNA in the global compartment. Thus, there are low amounts of MID1:RNA that could phosphorylate S6K in both compartments, leading to low values of the Hill function. From *q* = 20 to *q* = 23, both compartments show an increase in phosphorylated S6K once the threshold of the *l* function is crossed, but the effect is asymmetric. In the global compartment, [S6K^P^] rises steeply through the nonlinear part of the Hill curve, producing a strong boost. In contrast, the cluster compartment changes only modestly. This is because (i) clustering can only occur after MID1 has already bound RNA, and (ii) the clustering rate (*c*_8_ · *l*) is generally slower than the binding rate (*c*_1_ · *l*), limiting the overall impact. This change drives the sharp increase in total translation after *q* = 20.

**Figure S7:**
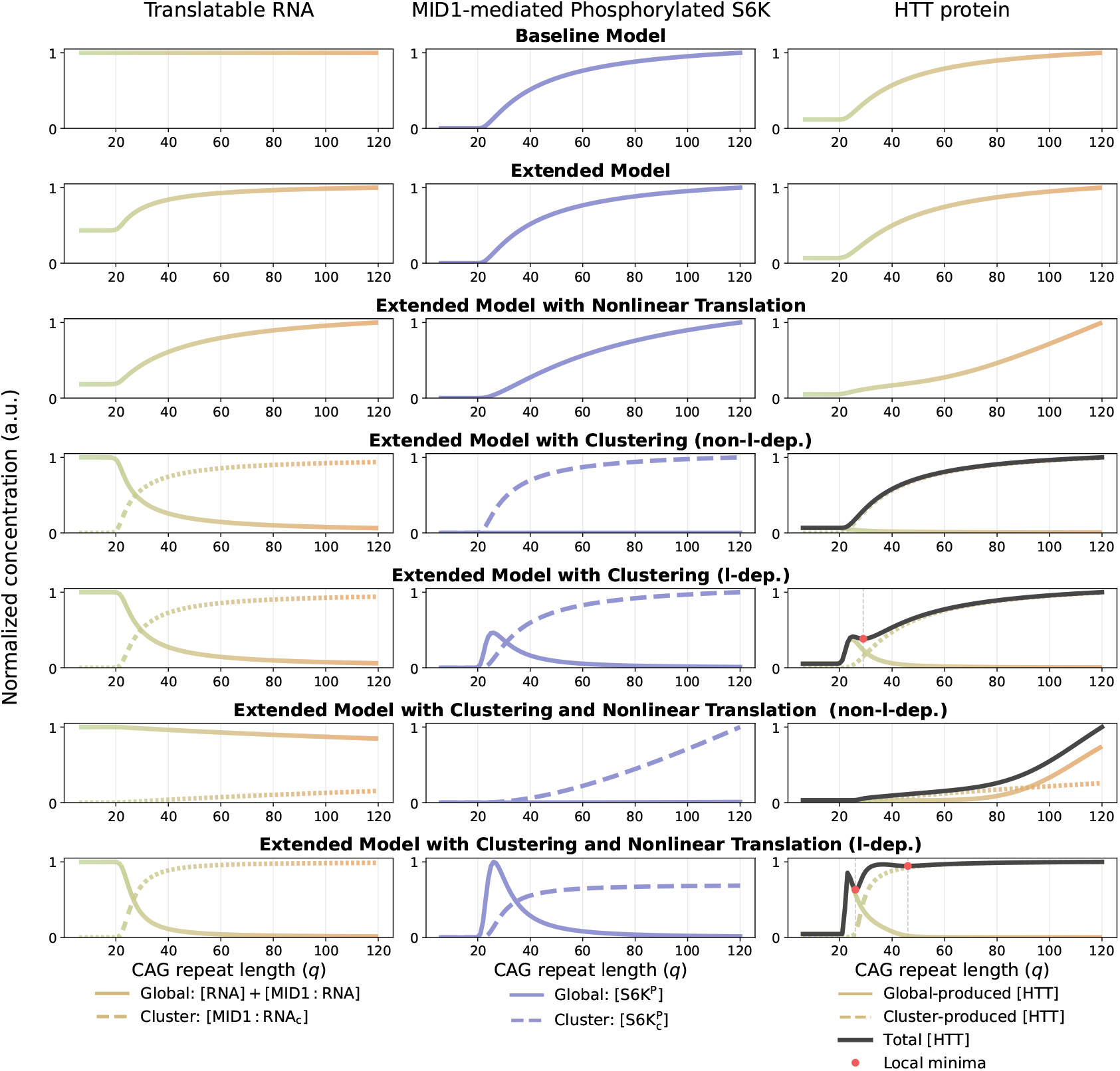
Simulation studies for all models with varying CAG repeat length. Steady state simulations for *q*_1_ = *q*_2_ ∈ [6, 120]: left, translatable RNA in global and cluster pools; middle, phosphorylated S6K in both pools; right, HTT levels in each pool and in total with local minima indicated.

Between *q* = 36 and *q* = 46, the cluster compartment is already in the saturation regime of its Hill curve, so translation increases only modestly. In contrast, the global compartment moves further down the declining [S6K^P^] regime, dropping below the midpoint of its Hill curve and causing a steep translation loss. This interplay explains the secondary dip in total translation described in the main text.

**Figure S8:**
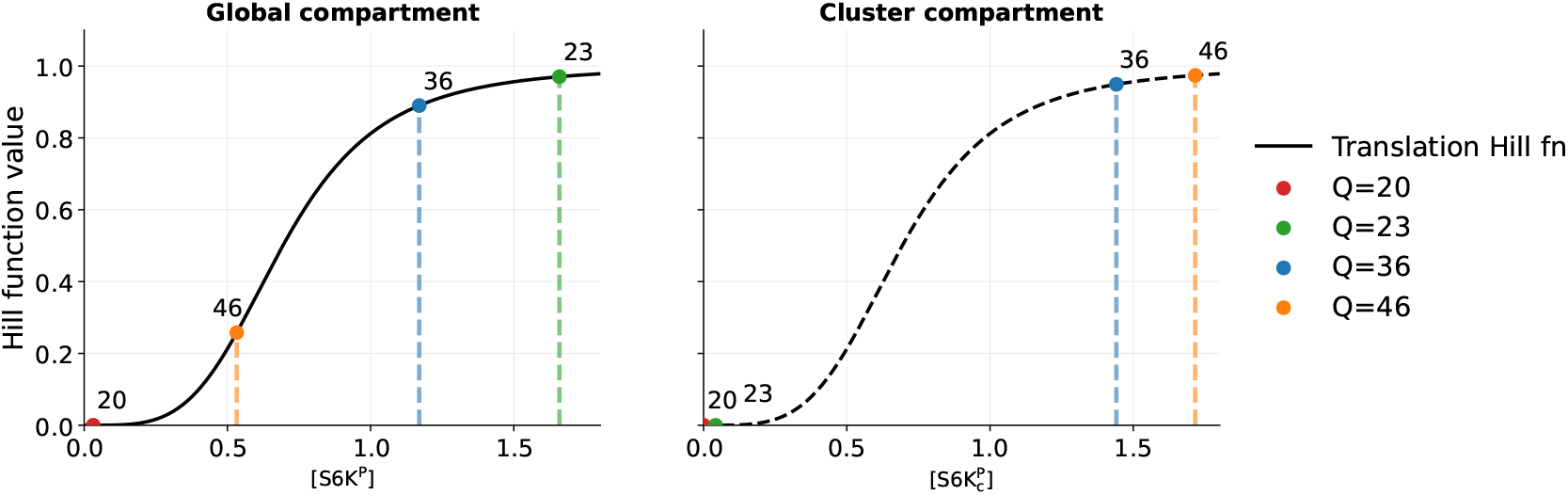
Effect of the Hill function on translation dynamics across CAG repeat lengths. Hill function output for global and cluster compartments at *q* = 20 (low translation in both), *q* = 23 (sharp post-threshold increase), *q* = 36 (peak after first drop), and *q* = 46 (second local minimum).

In the *Extended Model with Clustering* (l-dependent), the simulation results resemble those of the *Extended Model with Clustering and Nonlinear Translation* (l-dependent) (Fig. S7), with one key difference: the former lacks the second local minimum. This distinction arises because translation in the clustering-only model is governed by a linear rate function, which produces smoother dynamics. By contrast, the Hill-type function in the nonlinear translation model introduces threshold and saturation effects, allowing additional inflection points and thus the appearance of a second local minimum.

Overall, this analysis confirms that the shape of the Hill curve drives distinct phases of translation dynamics across CAG repeat lengths, resulting in the non-monotonic trends of the two models mentioned above. At the same time, these patterns may partly reflect artifacts of the current model formulation, and additional experimental data will be essential to confirm or refute them.

## S11 Noise model and residual analysis

As the data were obtained from five independent Western blot gels, gel-specific variabilityarising from differences in experimental conditions, sample handling, or imaging sensitivitymust be considered. To account for such variability, we assume five distinct noise parameters (*σ*_1_, …, *σ*_5_), one for each gel, so that every dataset is modeled with its own measurement noise distribution. This approach avoids the bias that would result from imposing a single global noise level across all experiments. The trade-off is increased model complexity, which can aggravate identifiability or overfitting, and in principle, too many noise parameters could distort the likelihood landscape if residuals reached zero (since *σ* → 0 would make it diverge), making optimization towards the optimal point difficult. In our case, however, multiple replicates that are different from each other prevent this situation, as the residuals would never be zero.

**Figure S9:**
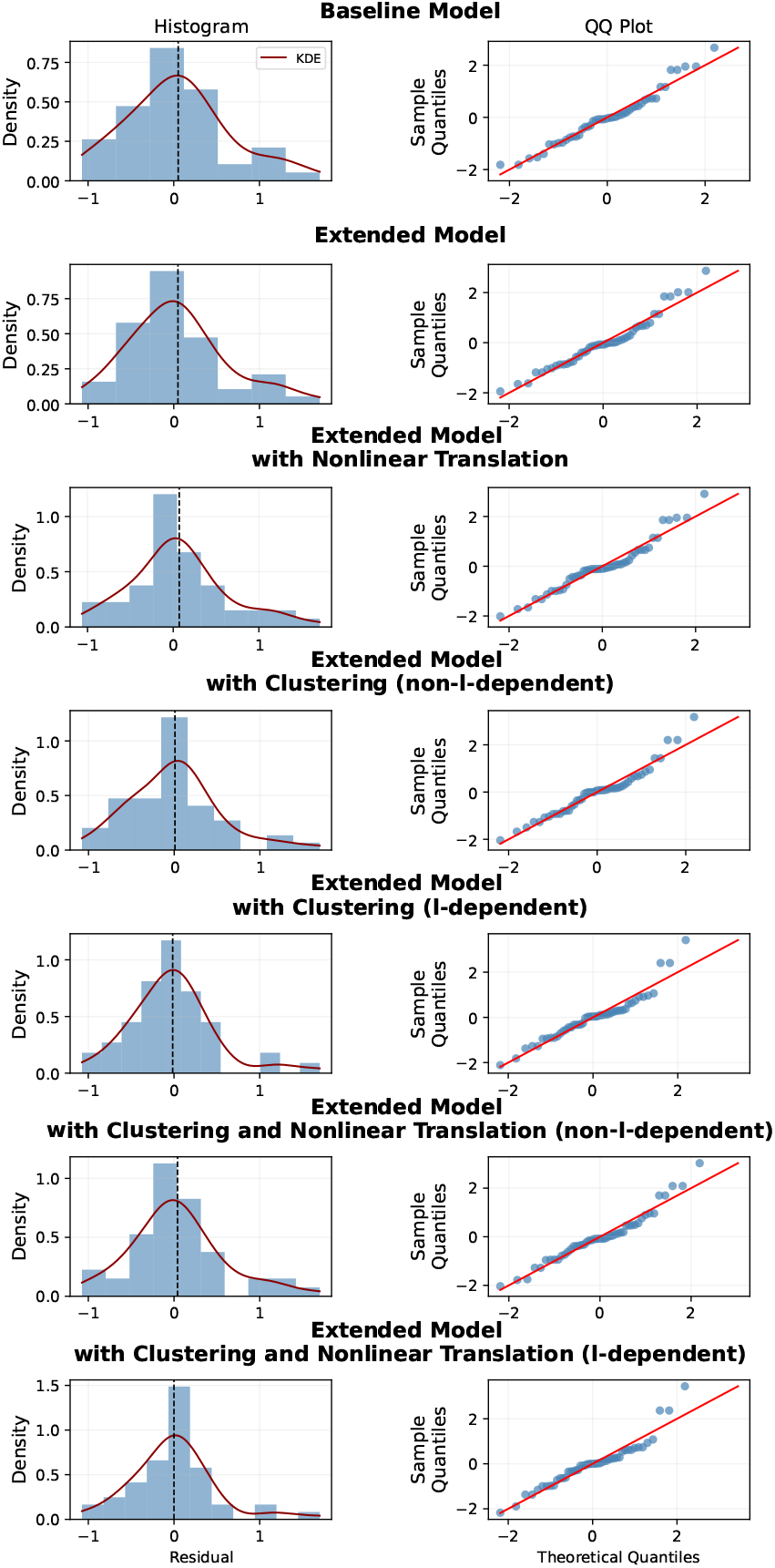
Residual diagnostics for all models. Histograms with kernel density estimates (left) and Q–Q plots (middle) are shown for each model. Improvements in normality and reduction of systematic residual patterns are most apparent in the extended models with clustering and nonlinear translation.

To assess whether the assumption of gel-specific noise parameters is reasonable, we examined residual diagnostics across all models (Fig. S9). The histograms, Q–Q plots, and residualversus-index plots show that residuals are approximately centered, with no major systematic biases. While the simpler models leave heavier tails and visible structure, the extended models with clustering and nonlinear translation yield more symmetric, near-normal residuals with reduced patterns. This supports the adequacy of the adopted noise model and suggests that our inference framework can capture the main sources of variability in the data.

**Figure S10:**
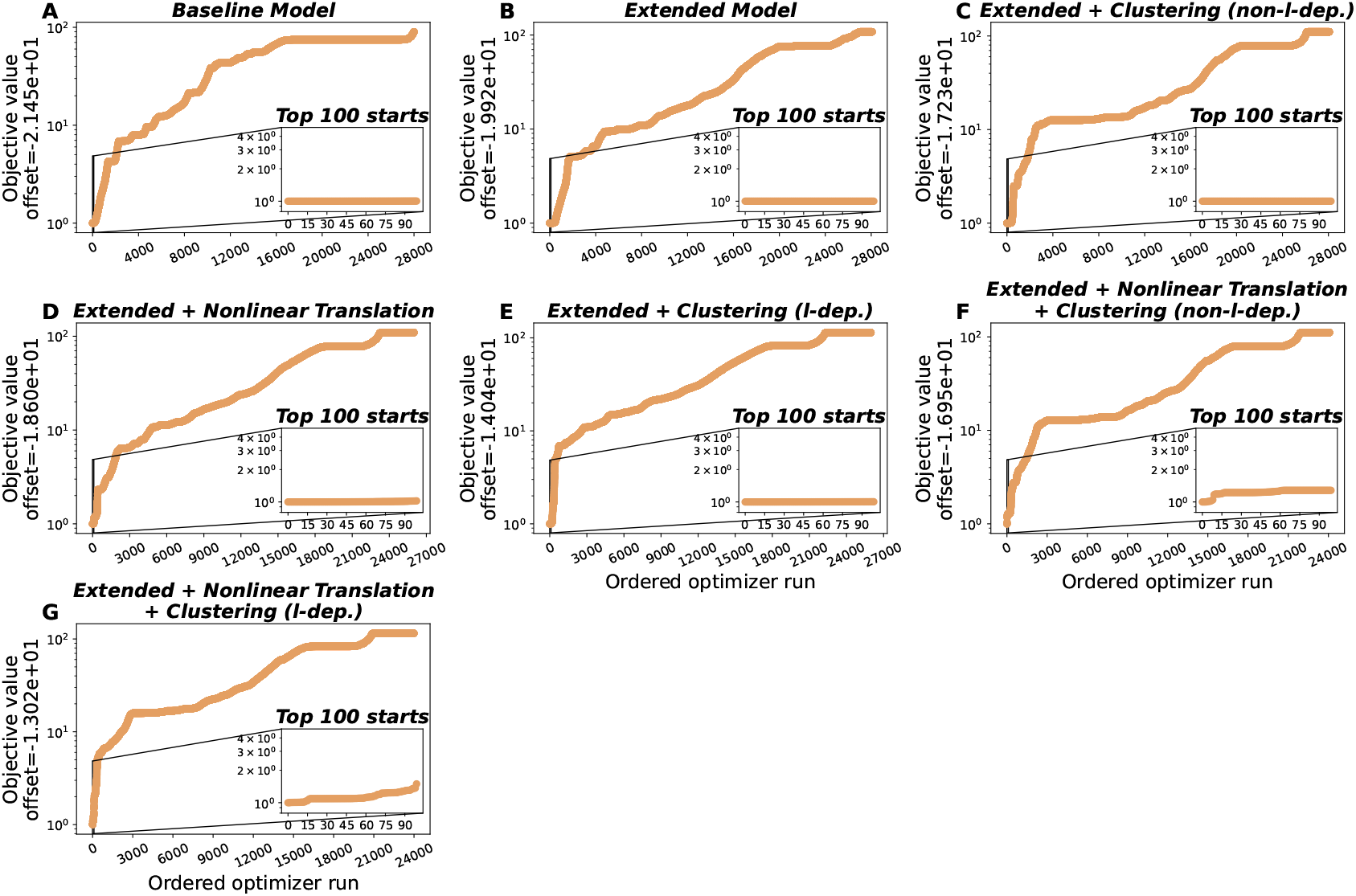
Waterfall plots for all seven models considered in this study. Each curve shows the negative log-likelihood (NLLH) values from independent optimization runs, sorted from best to worst. The *y*-axis is shown in log scale and offset by the best fit value, such that the minimum appears at 1; the applied offsets are indicated in the *y*-axis labels. Insets display the top 100 runs, with the *y*-axis scaled to the 95% confidence region (corresponding to an NLLH increase of 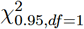 from the best fit). Rapid plateauing near the minimum indicates frequent convergence to the same optimum.

## S12 Convergence of optimization

To assess convergence in parameter estimation, we generated waterfall plots for all models considered (Fig. S10). Each plot shows the Negative Log-likelihood (NLLH) values obtained from independent optimization runs, sorted from best to worst.

Ideally, for well-constrained models with sufficient data, the objective function values from the best-performing optimization starts would rapidly plateau, indicating that the optimizer frequently converges to the same—or nearly the same—minimum. For more complicated models, the top ~100 starts do not always show just one plateau. Given the size and complexity of these models, coupled with the relatively limited amount of available experimental data, small differences in objective values among the top fits likely arise from flat directions in parameter space. In such cases, multiple parameter combinations can yield similarly good fits, but with slight numerical variation due to the optimizer exploring these nearly equivalent regions.

